# Using structure prediction of negative sense RNA virus nucleoproteins to assess evolutionary relationships

**DOI:** 10.1101/2024.02.16.580771

**Authors:** Kimberly R. Sabsay, Aartjan J.W. te Velthuis

## Abstract

Negative sense RNA viruses (NSV) include some of the most detrimental human pathogens, including the influenza, Ebola and measles viruses. NSV genomes consist of one or multiple single-stranded RNA molecules that are encapsidated into one or more ribonucleoprotein (RNP) complexes. These RNPs consist of viral RNA, a viral RNA polymerase, and many copies of the viral nucleoprotein (NP). Current evolutionary relationships within the NSV phylum are based on alignment of conserved RNA-directed RNA polymerase (RdRp) domain amino acid sequences. However, the RdRp domain-based phylogeny does not address whether NP, the other core protein in the NSV genome, evolved along the same trajectory or whether several RdRp-NP pairs evolved through convergent evolution in the segmented and non-segmented NSV genomes architectures. Addressing how NP and the RdRp domain evolved may help us better understand NSV diversity. Since NP sequences are too short to infer robust phylogenetic relationships, we here used experimentally-obtained and AlphaFold 2.0-predicted NP structures to probe whether evolutionary relationships can be estimated using NSV NP sequences. Following flexible structure alignments of modeled structures, we find that the structural homology of the NSV NPs reveals phylogenetic clusters that are consistent with RdRp-based clustering. In addition, we were able to assign viruses for which RdRp sequences are currently missing to phylogenetic clusters based on the available NP sequence. Both our RdRp-based and NP-based relationships deviate from the current NSV classification of the segmented *Naedrevirales*, which cluster with the other segmented NSVs in our analysis. Overall, our results suggest that the NSV RdRp and NP genes largely evolved along similar trajectories and that even short pieces of genetic, protein-coding information can be used to infer evolutionary relationships, potentially making metagenomic analyses more valuable.

## Introduction

The International Committee on Taxonomy of Viruses (ICTV) established two phyla for RNA viruses with a negative sense viral genome: the *Ribozyviria* and *Negarnaviricota*. Many important human RNA viruses are currently classified among the *Negarnaviricota*, including the influenza A virus (IAV), Rift Valley fever virus (RVFV), Lassa virus (LASV), measles virus (MeV), rabies virus (RABV), and Ebola virus (EBOV). We will therefore only consider the *Negarnaviricota* here and refer to them as negative sense RNA viruses (NSV) for simplicity (**Figure 1)**. The genomes of NSVs consist of single-stranded, negative sense RNA that is copied in the context of a ribonucleoprotein (RNP) complex. Each RNP is composed of a viral RNA (vRNA) template, an RNA polymerase containing an RNA-dependent RNA polymerase (RdRp) domain, and numerous nucleoproteins or nucleocapsids (NP) (1, 2). The self-oligomerizing NP molecules make up the majority of each RNP, and they protect the genome from degradation, act as scaffold for RNA structure formation, and assist in viral replication by acting as a processivity factor (3, 4).

**Figure 1:**
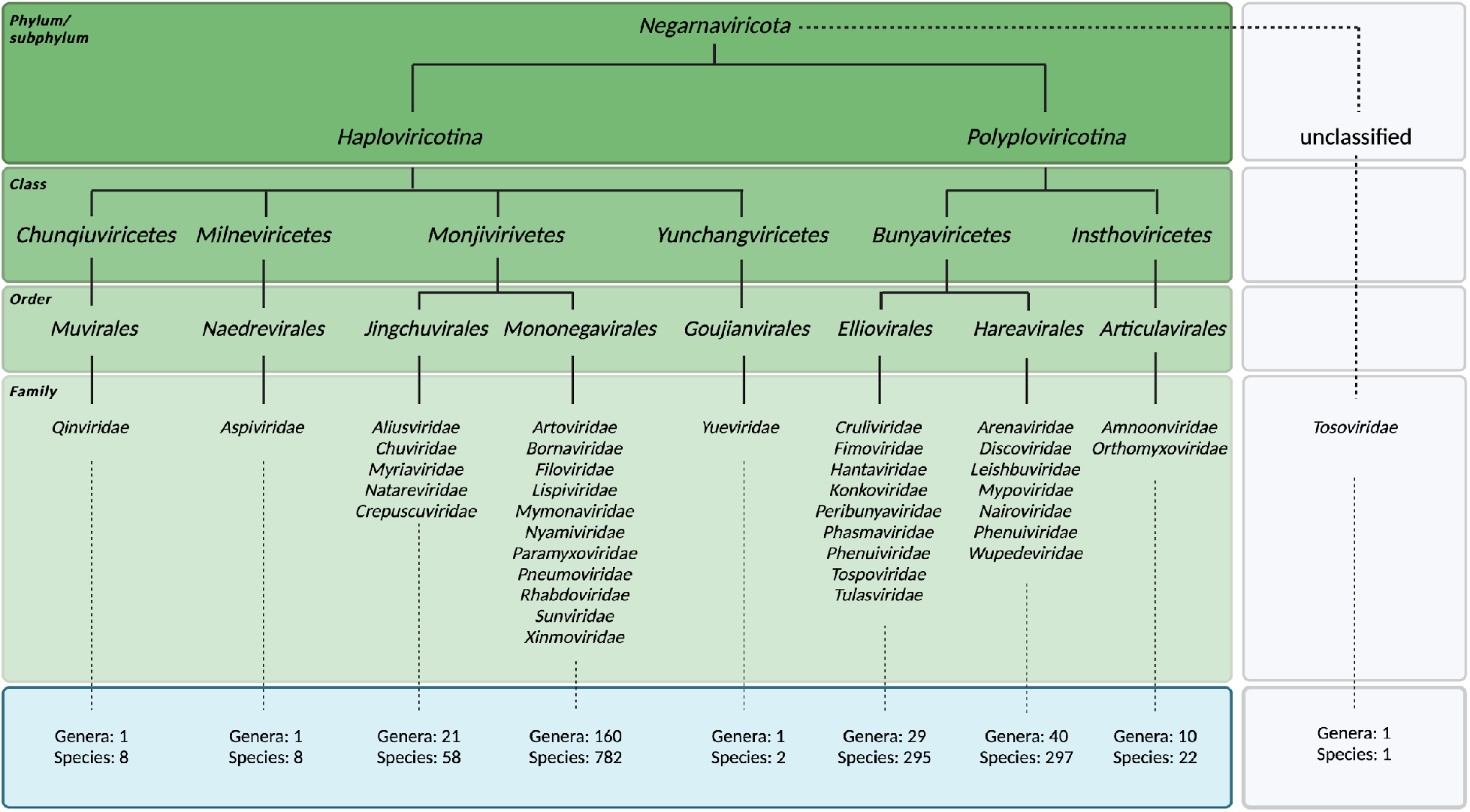
Classified viral species within *Negarnaviricota*. Taxonomic breakdown of the *Negarnaviricota* phylum of the *Riboviria* realm according to the ICTV and NCBI databases at the time of manuscript revision. The phylum consists of all NSVs and is divided into the subphyla *Haploviricotina* and *Polyploviricotina*. Most classified viruses either show presence or absence of mRNA capping activity in the RNA polymerase and the organization (non-segmented or segmented) of the viral genome, depending on whether they are assigned to the *Haploviricotina* or *Polyploviricotina*, respectively. Note that the *Naedrevirales* have a segmented genome, but an RNA polymerase with mRNA capping activity. Additional exceptions are listed in the introduction.

Despite these shared features, both the genome organization, and replication and transcription mechanisms vary widely among NSVs (5). A key division between NSVs can be made based on the presence or absence of genome segmentation. In the absence of genome segmentation, all NSV genes are encoded on the same RNA molecule and transcribed in sequence. By contrast, NSVs that have segmented genomes encode key viral genes such as the RNA polymerase and NP on separate RNA molecules. In the case of IAV, the RNA polymerase is even divided up into three subunits that are each encoded by a separate RNA molecule. A further division between NSVs can be made based on the molecular mechanism that they use to initiate transcription of the genome or genome segments. It is presently not fully understood how the different NSV genome architectures and RNA synthesis mechanisms evolved.

Every virus within the *Negarnaviricota* phylum shares a common two-gene core: the RNA polymerase gene (or the three genes encoding its subunits) and an NP gene (6). The RdRp domain of the large protein (L) - or the PB1 subunit of the heterotrimeric IAV RNA polymerase - is approximately 300 amino acids long and conserved amongst all RNA viruses (kingdom *Orthonavirae*), whereas the rest of the RNA polymerase is not. As the core proteins of the RNP, the RNA polymerase and NP molecules need to work together to replicate and transcribe the viral genome, and their functions must work in concert with the different genome organizations. It is possible that different NSV RdRp-NP pairs diverged from a single common ancestor RdRp-NP pair, and that their evolution is linked to each other as well as the genome architecture. Alternatively, different RdRp-NP pairs may have evolved through convergent evolution, potentially as a result of changing (and reassorting or recombining) NSV genome architectures. Understanding the properties and evolutionary history of the RdRp domain and NPs may shed light on how different NP molecules work together with different RNA polymerase activities and support different NSV genome and RNP structures.

The conserved RdRp domain has been the focal point for many evolutionary analyses (7, 8). While traditional phylogenomics utilizes metagenomics data, sequence alignments, and comparison metrics, the ever-expanding wealth of structural information provides a potential avenue for the refinement of important branching points (9–17). In particular, structural comparisons of RdRp domains of different virus families have illustrated surprising similarity amongst *Orthomyxoviridae* (with negative-sense ssRNA), *Flaviviridae* (with positive-sense ssRNA) and *Cystoviridea* (with dsRNA), suggesting NSVs originated from dsRNA viruses which in turn evolved from positive-sense ssRNA viruses (14). These phylogenies combined with analyses by the ICTV have mapped out six major phyla of *Orthonavirae* and elucidated ancestral trends amongst them. NSVs subsequently evolved into the two subphyla *Haploviricotina* and *Polyploviricotina* (**Figure 1**) (18).

The division of NSVs into two subphyla separates NSVs that have an RNA polymerase that possess mRNA capping activity from NSVs that have an RNA polymerase that performs cap-snatching to generate capped viral mRNAs, respectively (8). Additionally, the majority of NSVs within the *Haploviricotina* subphylum contain continuous non-segmented genomes, while the majority of viruses in the *Polyploviricotina* subphylum contain segmented genomes. Genome segmentation allows for reassortment of gene segments, which can contribute to the emergence of pandemic viruses and the spread of immune or antiviral resistance mutations (19, 20). Segmentation may also underlie mechanisms of viral differential gene expression (21, 22). On the other hand, segmentation complicates single-partite virus genome packaging. A virion of a segmented single-partite virus is not viable unless it contains the entire set of viral genome segments. The coordination and packaging of all segments is thus a crucial step in segmented, single-partite NSV infections. In IAVs, the majority of virion particles contain one copy of each of the eight genome segments (23).

Exceptions to the above apparent rules of RNA polymerase function and segmentation have been identified. These exceptions include viruses classified in the order of the *Naedrevirales*, which encode an RNA polymerase with mRNA capping activity but have a segmented genome, viruses in the genus *Dichorhavirus* of family *Rhabdoviridae*, which similarly have a segmented genome but an RNA polymerase homologous to members of the *Haploviricotina*, and viruses in the *Chuviridae* family, which have many different genome organizations, including non-segmented and segmented genomes. A final exception that we will mention are the *Tulasviridae*, a family of ambisense RNA viruses that are classified within the *Bunyavirales* order, but that have non-segmented genomes. When additional NSVs are discovered in the future, further exceptions to the capping and segmentation-based classification may be found.

As mentioned above, previous work has used analyses of RdRp domain sequences to understand the evolution of genome structure and segmentation within *Negarnaviricota* or the relations among NSV subphyla, classes, and orders (6, 24). Since the RNA polymerase works together with NP to replicate and transcribe the viral genome, studying the evolutionary history of NPs may reveal additional insights into the evolution of NSV genome replication as well as its segmentation. However, NP sequences are too short for robust phylogenetic analyses. We here combined experimentally obtained structural data and the deep learning tool AlphaFold 2.0 (AlphaFold2) (**Figure 2**) to explore if the evolutionary relationships among NSVs can be reconstructed from NP sequences and if this relationship provides additional insight into the divergence of the NSV subphyla and the emergence NSV genome segmentation. We find that the structural homology of NPs within NSVs provides more phylogenetic information than NP sequences alone. In addition, we observe that the NP structural clustering largely matches the clustering of the RdRp domain, suggesting that the two NSV core genes likely co-evolved. We suggest that our approach may be useful for metagenomics and virus discovery studies where RdRp sequences may not be complete or missing.

**Figure 2:**
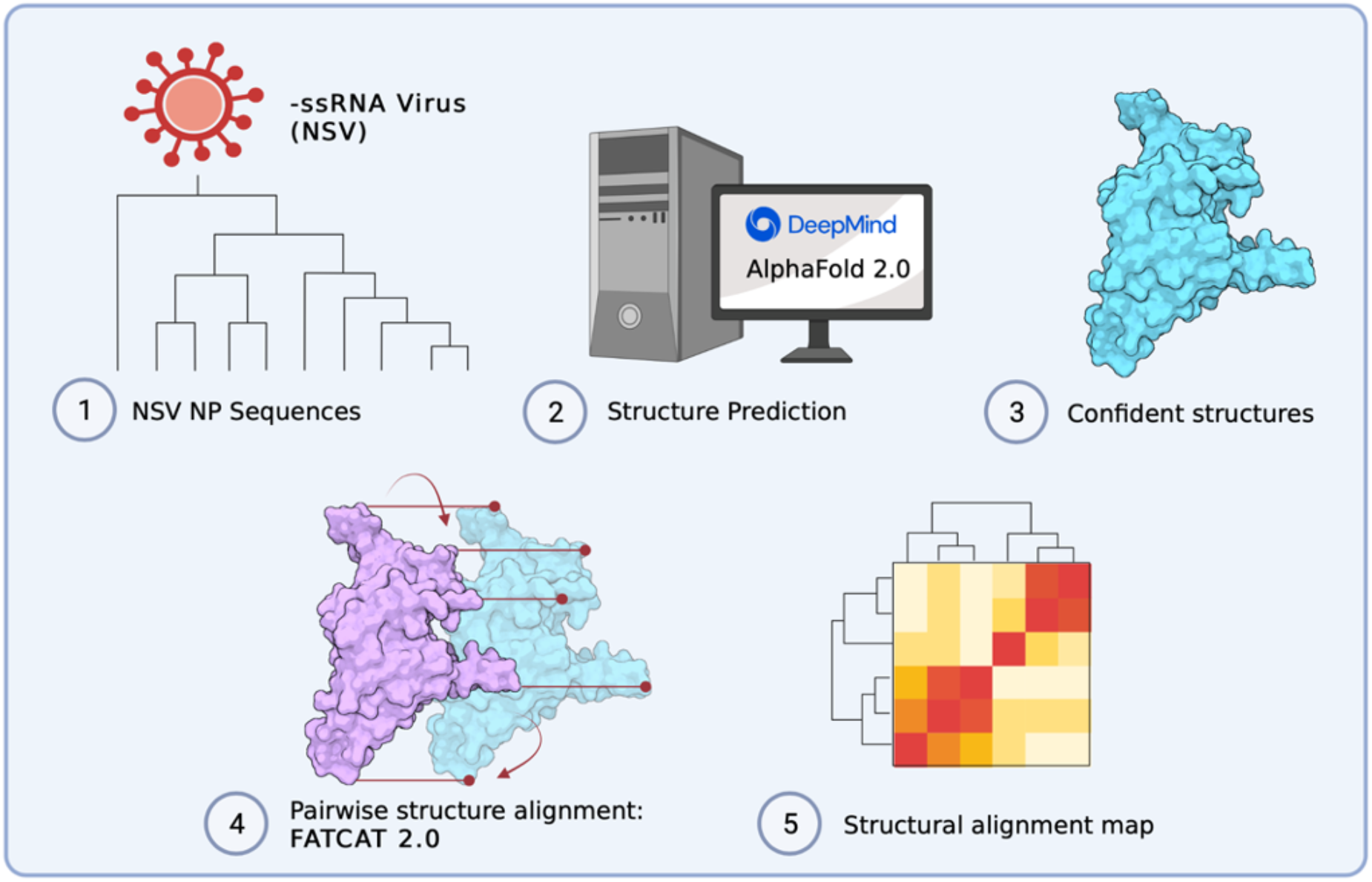
Overview of computational approach to build structural alignment map. (**1-2**) Data was acquired based on sequence availability. AlphaFold 2.0 was used to predict structures. (**3-4**) AlphaPickle was used to visualize confidence metrics, following which structures were selected based on confidence score cut-offs and individual pairwise flexible alignments were computed using the FATCAT 2.0 package compiled on a high performance cluster (HPC). (**5**) Data processing was performed to parse structural alignment data into a concise heatmap for comparison with MSA percent identity matrices.

## Results

### Experimental NP structure dataset

At the time of revision of this manuscript, there were 858 viral species classified as members of the *Haploviricotina* subphylum and 614 species assigned to the *Polyploviricotina* subphylum in the ICTV database. The taxonomical breakdown is summarized in **Figure 1**. NP structural data was found for 34 of these NSVs in the PDB database. Within this initial dataset, 21 NP structures were from the *Polyploviricotina* subphylum and 13 from the *Haploviricotina* subphylum. Given the relatively even ratio of currently classified viruses in each subphylum in the ICTV, this initial NP structure dataset appears to be biased towards the human-disease causing, segmented NSVs (**Figure 3**).

**Figure 3:**
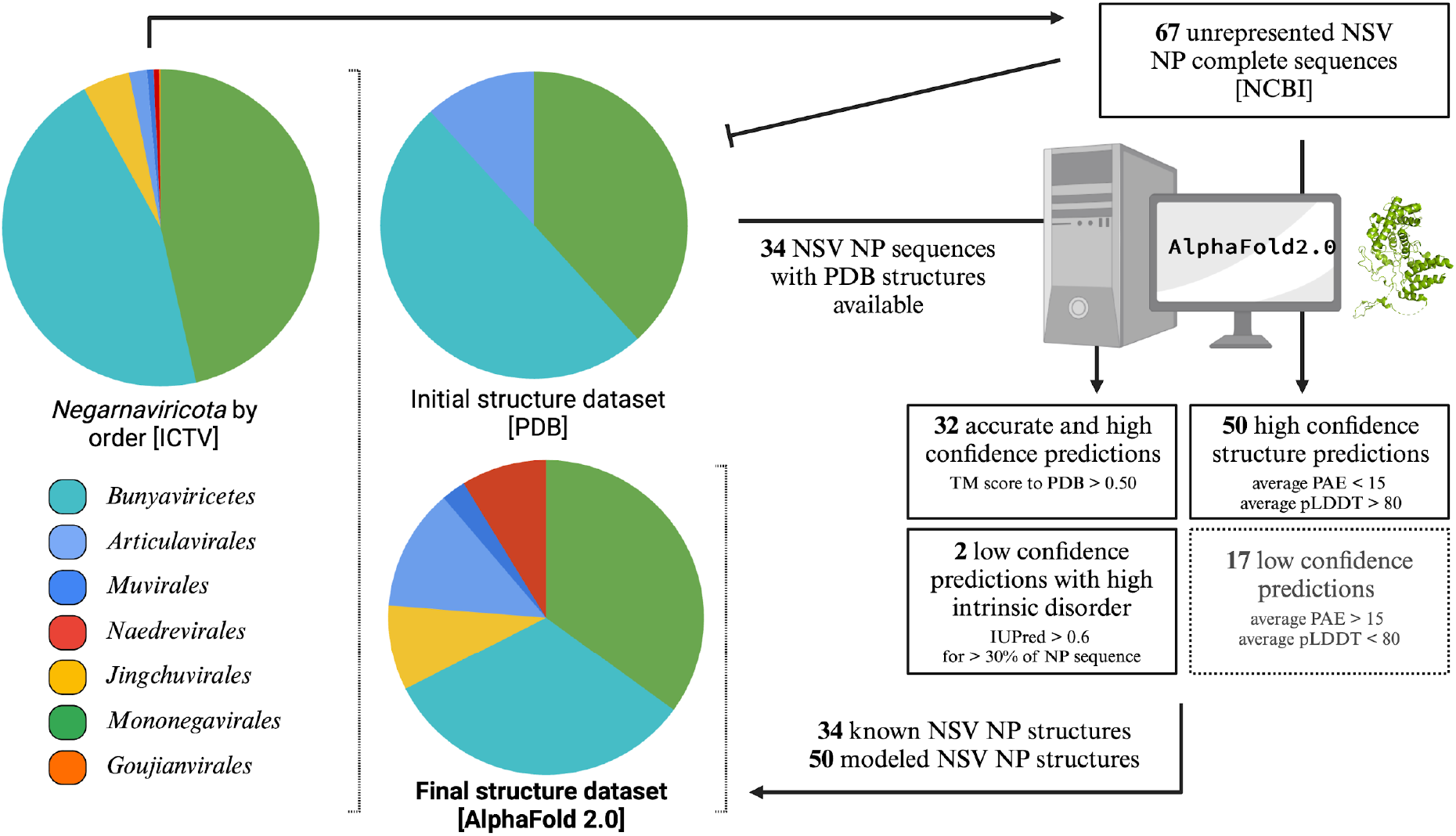
Composition of the initial and final structural dataset of *Negarnaviricota* NPs. The composition breakdown of *Negarnaviricota* by phylogenetic order (*left*) was determined by the number of species in each order in the ICTV/NCBI database at the time of the manuscript preparation. The order distribution in percentages is as follows: 45.6% *Bunyaviricetes*, 1.8% *Articulavirales*, 0.6% *Muvirales*, 0.6% *Naedrevirales*, 4.8% *Jingchuvirales*, 46.4% *Mononegavirales*, and 0.2% *Goujianvirales*. At the time of manuscript revision, there were 34 NSV NP structures available in the PDB database with representation from only the *Bunyavirales, Articulavirales*, and *Mononegavirales*. The final NSV NP dataset included a total of 84 structures generated with AlphaFold 2.0.

### NP structure prediction using AlphaFold2

Phylogenetic analyses can be performed on many types of information, including multiple-sequence alignments and structural alignments. We wondered if the current NP structure dataset could be expanded using structural prediction methods like AlphaFold2. To explore this question, we first determined if AlphaFold2 could accurately predict the structures of the known NPs based on the primary sequences of the 34 NP structures in the PDB. To minimize the impact of the known NP structure on the prediction, the AlphaFold2 settings were restricted to structure templates released prior to the release dates of the NP structures in the PDB. As shown in Fig. 4A, the overall performance of AlphaFold2 was robust, with 27 out of 34 predictions having high overall model confidence (average pLDDT scores above 70) (***Supplementary raw data file 1***). Aligning the predicted structures to the corresponding experimental structures produces TM-scores above the 0.50 threshold for 30 out of 34 structures, suggesting that the predictions are in fact of the same general topology as the experimental structures (25).

**Figure 4:**
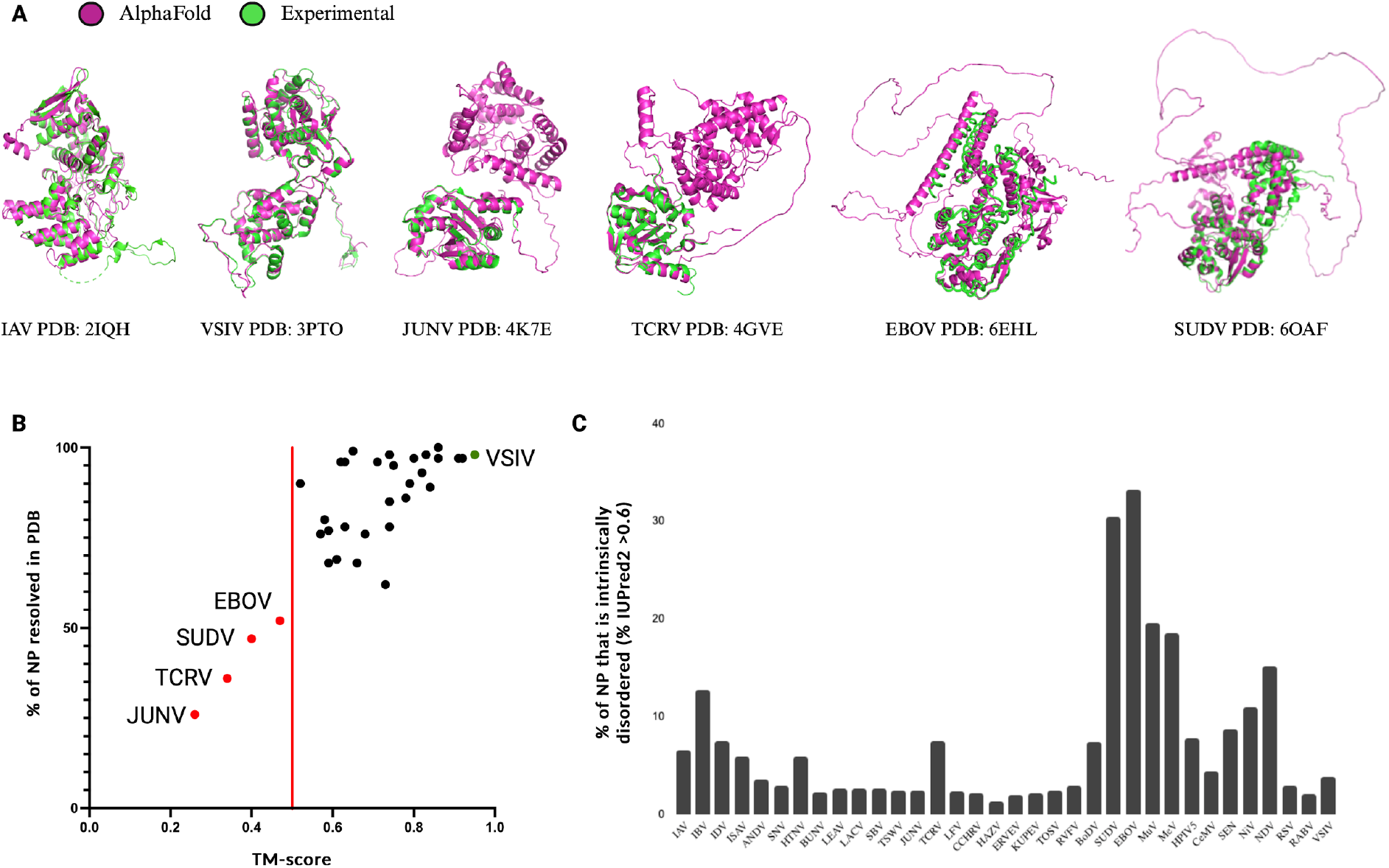
AlphaFold2 structure prediction performance on NSV nucleoproteins in the PDB. Overall prediction performance of AlphaFold2 on NSV NPs was assessed using an initial dataset of 34 NSVs with (partial) experimentally solved structures deposited in PDB. (**A**) Structural alignments of AlphaFold2 predictions (magenta) and experimental structures (green) are displayed for the most accurate prediction (VSIV and IAV, left), a partially resolved experimental structure with the lowest % of NP modeled (JUNV and TCRV, middle), and a highly intrinsically disordered structure (SUDV and EBOV, right). (**B**) The predicted AlphaFold2 structures were compared to the experimental NP structures using rigid jFATCAT structural alignment. Next the percentage of NP residues that was experimentally resolved and deposited in the PDB was plotted against the alignment score (TM-score). Four structures fell below a TM-score of 0.50 (red line) and are depicted as red points and labeled. The top scoring AlphaFold2 prediction with an almost perfect alignment (VSIV NP) is denoted by the green data point. (**C**) Intrinsic protein disorder predictions for the initial dataset using IUPred2 show a high percentage of disorder within the SUDV and EBOV NPs.

Next, we aligned the AlphaFold2 structures with the corresponding experimental structures using jFATCAT. Overall, we find good agreement between the experimental and computed structures, with the VSIV NP showing the best agreement with a TM-score of 0.95 (**Figure 4A**). However, for some NPs we observed deviations in regions of computed intrinsic disorder, which many experimental structure-solution methods were unable to resolve (**Figure 4A**). Only four structures seemed to have a low resemblance to the experimental structure, with TM-scores less than 0.50 (**Figure 4B**). Further analysis showed that the TM-score of the AlphaFold2 structure correlated with the percentage of modeled NP residues and/or the percentage of sequence coverage of the experimental structure (**Figure 4B**). Thus, the experimental NP structures with limited sequence coverage or missing residues (i.e., the Junin arenavirus (JUNV), Tacaribe virus (TCRV), Sudan virus (SUDV), and Ebola virus (EBOV) NPs) align poorly with the AlphaFold2 predictions of the complete NP sequences.

Investigating the four, relatively poorly aligned NP models in more detail, we find that both the experimental JUNV NP structure (PDB: 4K7E) and the TCRV NP structure (PDB: 4GVE) contain only the C-terminal domains (**Figure 4A**), while the AlphaFold2 models contain both C-terminal and N-terminal domains. Superposing the partial experimental NP structures from PDB with the AlphaFold2 models shows good alignment between the C-terminal domains (**Figure 4A**). The remaining two outliers, EBOV NP (PDB: 6EHL) and SUDV NP (PDB: 6OAF), have experimental structures of which only the N-terminal core domain is resolved. The unresolved C-terminal domains of both the EBOV and SUDV NPs are predicted to be intrinsically disordered domains. The AlphaFold2-predicted structures align with the N-terminal core domains that are resolved in the experimental structures, while the parts of the AlphaFold2-predicted structure that do not align contain large unstructured loops that have low prediction confidence (**Figure 4A**). The AlphaFold2 predictions follow the IUPred2 protein disorder prediction for the sequences in their entirety (26) (**Figure 4C**). The unstructured regions and low-confidence loops in the AlphaFold2 predictions correlate with the regions of high disorder prediction (regions longer than a few amino acids with disorder prediction scores above 0.50). Analysis of the predicted disordered regions in the 34 NP models shows that large regions of intrinsic disorder (>20%) are uniquely present in the *Filoviridae* NPs (**Figure 4C**). These disordered regions are likely inherent properties of individual NPs and not an erroneous artifact of the AlphaFold2 analysis. We thus consider the AlphaFold2 predictions sufficiently robust for the majority of NP structures to expand the experimental NP structure dataset.

### Expanding the NP structure dataset using AlphaFold2

We next used AlphaFold2 to extend the experimental NP structure dataset to obtain better structural coverage of the *Negarnaviricota* NPs. To this end, additional viral species with known and complete NP sequences were chosen from unrepresented NSV families. Next, the NP structures of these viruses were predicted using AlphaFold2. The structure predictions that passed the confidence requirements (average PAE scores <15 and average pLDDT scores >80) were included in the final NP structure dataset (**Figure 3**). Please see **Supplemental Information** for data on the structure predictions that were excluded. The expanded final dataset contained a total of 84 NP structures with 57.2% from *Haploviricotina* and 42.8% from *Polyploviricotina* (**Supplementary Table 1**).

### Analysis of the NP and RdRp domain MSAs

To assess if the NP structures contained enough information to investigate the evolutionary history of NP, we first generated an MSA of the RdRp domain sequences to use as a benchmark. Only 82 of the 84 NSVs were included in this MSA as the freesia sneak virus (FSnV) and tulip mild mottle virus (TMmV) RdRp sequences were not available in sequence databases at the time of analysis. The MSA rows were arranged as shown in **Supplemental Table 1** and the resulting MSA was used to generate a percent identity matrix. This matrix is shown as a heatmap in **Figure 5, left**. We next generated an MSA using the 84 NP sequences for each NSV and illustrated the NP percent identity matrix as a heatmap in the same order as the RdRp MSA heatmap (inclusive of FSnV and TMmV) (**Figure 5, right**).

**Figure 5:**
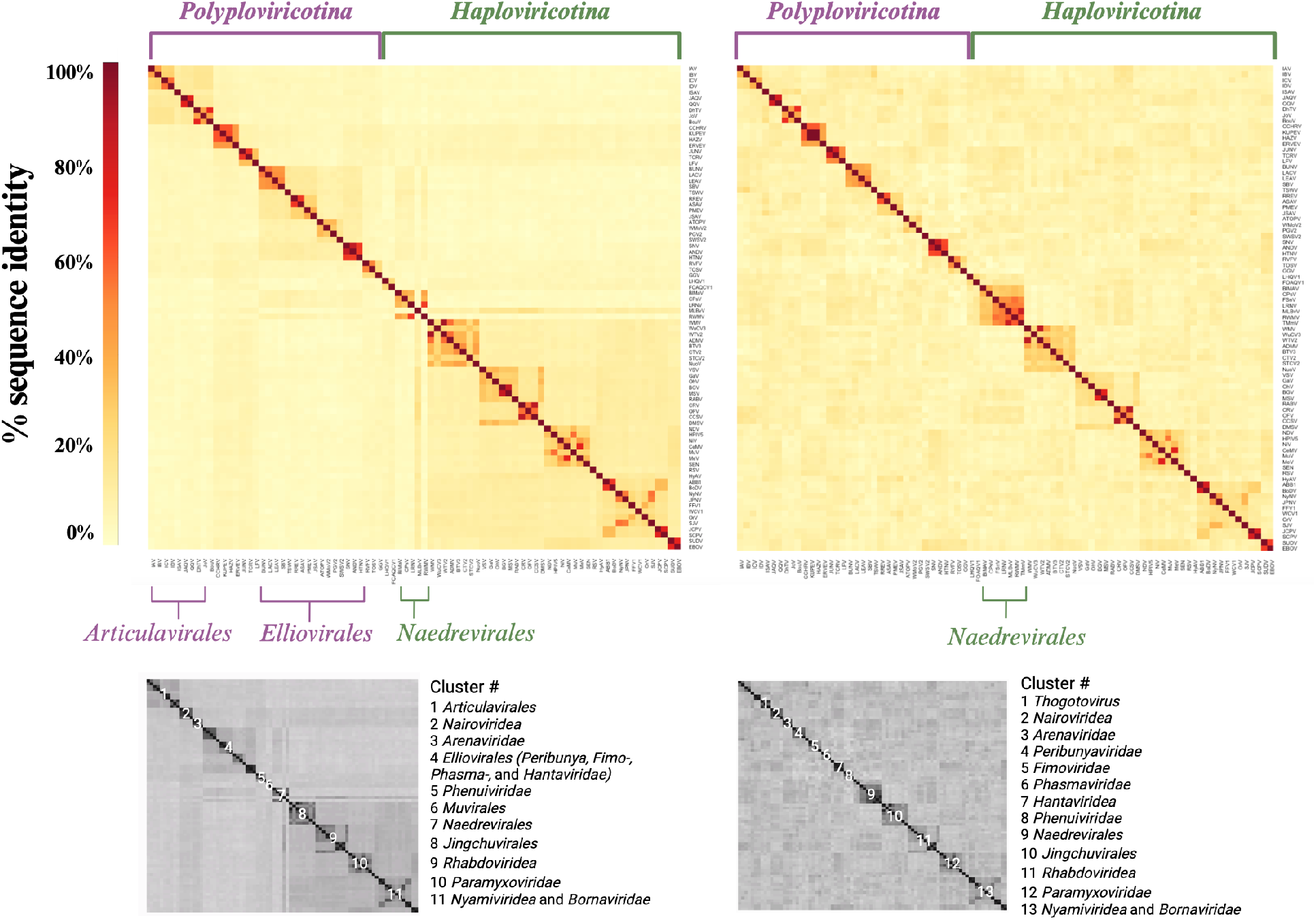
NSV RdRp multiple sequence alignment heatmap shows distinct clustering not seen with NP. RdRp (*left*) and NP (*right*) amino acid sequence MSAs illustrated as percent identity matrices in a fixed order based on current phylogeny. The RdRp MSA does not include 2 of the 84 NSVs used in the NP MSA, as complete freesia sneak virus (FSnV) and tulip mild mottle virus (TMmV) RdRp sequences were not available in sequence databases at the time of analysis. The two subphyla are indicated as well as the resolved virus order and family clusters.

The RdRp and NP heatmaps shown in **Figure 5** are presented with an identical x- and y-axis order, with the exception of FSnV and TMmV, which are absent in the RdRp MSA heatmap, due to incomplete sequence availability at the time of the analysis. The clustering can thus be qualitatively compared across the two sequence alignments. The predefined order of the heatmaps separates the *Polyploviricotina NSVs* (mostly segmented) annotated in purple from the *Haploviricotina* NSVs (mostly non-segmented) annotated in green. The RdRp MSA heatmap shows four darker sub-clusters in the top left corner that are grouped into a single larger cluster. This larger group consists of 10 NSVs that all belong to the *Articulavirales* order, in line with the current ICTV classification. Comparing the RdRp and NP MSA heatmaps, we note that the virus order cluster is lost in the NP heatmap.

The second major cluster in the RdRp MSA heatmap represents the *Elliovirales* and includes several sub-clusters that represent the virus families including the *peribunyaviridea*, the *Tospoviridea*, the *Fimoviridae*, the *Phasmaviridae*, and the *Hantaviridae*. The *Hareavirales* are spread over three sub-clusters that represent the virus families *Nairoviridae, Arenaviridae* and the *Phenuiviridae*. Similar to the *Articulavirales* clusters in the RdRp MSA heatmap, the *Elliovirales* cluster is lost in the NP MSA heatmap, but the different NSV families readily form identifiable clusters (**Figure 5**).

Continuing down the diagonal to the right of the RdRp MSA heatmap, we enter the *Haploviricotina* subphylum. The first tiny cluster represents the *Muvirales* order. This cluster is followed by a disjointed cluster of five members of the *Naedrevirales*. In this cluster, mirafiori lettuce big vein virus (MLBvV) appears to be more similar to the subsequent *Jingchuvirales* order cluster than to the *Naedrevirales* cluster. In the NP MSA heatmap, two additional members of the *Naedrevirales* order are included (FSnV and TMmV) and they neatly cluster with the other order members, creating a sub-cluster of seven *Naedrevirales*. It is interesting that the available *Naedrevirales* RdRp sequences do not provide the same level of clustering as the NP sequences despite the established phylogenetic relationships that are based on the RdRp gene. It is important to note here that, at the time of analysis, the *Neadrevirales* order is taxonomically classified within *Haploviricotina* based on the mRNA capping ability of their encoded RNA polymerase, even though their genomes are segmented. The *Naedrevirales* cluster in the RdRp MSA heatmap appears to have more sequence homology to the *Elliovirales* cluster than to the large *Haploviricotina* cluster, suggesting that in spite of the capping domain, their NTP incorporation function may be more closely related to the *Elliovirales* order.

In the large *Haploviricotina* cluster, the eight members of the *Jingchuvirales* order form a cluster. This cluster is nearly identical in the RdRp and the NP MSA heatmaps. The remaining 31 NSVs belong to the Mononegavirales order, but they do not form a single cluster. Instead, we observe small clusters that represent their respective NSV families. The clearest clusters are formed by the *Rhabdovirideae*, the *Paramyxoviridae*, and the *Nyamiviridae*. The latter family is flanked by members of the *Bornaviridea* family, with which they form a larger cluster. These four clusters are present in the NP heatmap as well. The members of the *Pneumoviridae*, two *Qinviridae*, one *Lispiviridae*, and two *Filoviridae* in our dataset are part of the larger *Haploviricotina* cluster. In the NP MSA heatmap, this larger *Haploviricotina* cluster is not evident.

Overall, this sequence level analysis suggests that the virus family clusters are resolved in both the RdRp and NP heatmap, and that the family clustering is consistent with the current phylogeny of *Negarnaviricota* (27, 28). However, at a higher level, the RdRp MSA heatmap is able to identify some NSV orders and more distant homologies between family clusters that the NP MSA heatmap can not. Thus the RdRp sequence appears to offer more phylogenetic information to derive long-distance ancestry than the NP sequence.

### Pair-wise analysis of NP structures

We hypothesized that the NP AlphaFold2 structure dataset contained more phylogenetic information than the NP sequences alone, because protein structures are generally more conserved than the primary sequence. To explore this, we had to align our AlphaFold2 NP structures. Aligning structural information requires a different approach and significantly more computational power than aligning primary amino acid sequence data. Foundational structural alignments utilize rigid body backbone algorithms. While this is a powerful method for assessing the similarity of closely related structures that are solved in similar conditions, the natural motion and dynamics of protein structures are not accounted for in rigid body alignments (29, 30). As proteins can adopt multiple conformations according to their function or protein structures have different energy minima, it is critical to analyze structures in a way that tests whether two proteins are identical, even if they have adopted different conformations (*e.g*., “open” or “closed”, or “active” or “inactive”). Various complex algorithms have been developed to attempt to incorporate dynamic motion into structural alignments. For our NP AlphaFold2 structure analyses, we used the Flexible structure AlignmenT by Chaining Aligned fragment pairs allowing Twists (FATCAT) algorithm (31). This method utilizes fragment pairs and iterative rotations/twists to converge on an alignment that would account for alternate conformations. As is the case with NSV NPs, the two-domain NP structures have been proposed to twist upon oligomerization and/or RNA-binding to form the helical chains seen in RNP complexes (32–35).

Pairwise alignments for all possible pairs of NPs within the dataset were individually computed. Each alignment pair resulted in an alignment p-value, which we interpret as the probability of observing a more identical alignment between the structure pairs. A structural alignment heatmap was generated from the alignment p-values and clustered using the Ward hierarchical clustering method (**Figure 6**). The diagonal represents self alignment, or a p-value of exactly zero. It is important to note that the color scale of the FATCAT heatmap (**Figure 6**) is not directly comparable to the MSA heatmaps (**Figure 5**) as the values illustrated in each case derive from fundamentally different calculations.

**Figure 6:**
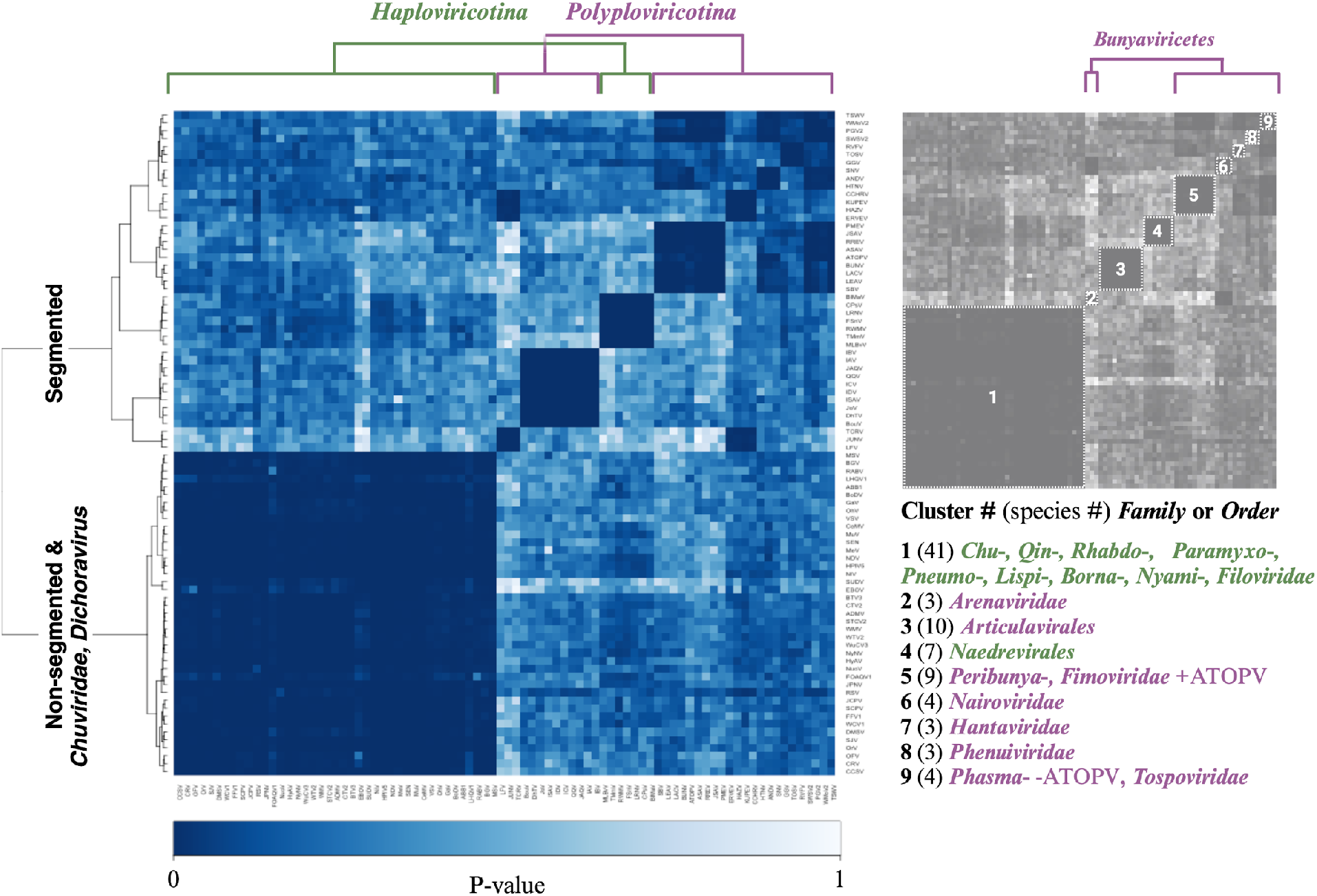
NSV NP FATCAT Alignment by Ward hierarchical clustering. FATCAT Pairwise structural alignment results for 84 NSV nucleoproteins represented as a heatmap of p-values. The data was clustered using the Ward hierarchical clustering method. The diagonal represents self alignment, or a p-value of exactly zero. Numbered clusters are defined in the key to the right of the heatmap. The number of virus species in each cluster is indicated in parentheses.

The pairwise flexible structure alignment data was clustered by hierarchical clustering using the Ward method. The resulting heatmap and dendrogram are presented in **Figure 6**. The dendrogram splits the AlphaFold2 NP structures into two main clusters and nine smaller clusters, which we identified and annotated in the key to the right of the heatmap. Starting from the left side of the heatmap, cluster 1 includes 41 of the 48 *Haploviricotina* NSVs in our dataset. The low p-values within the first cluster suggest a high structural homology among the virus families, which include the *Chuviridae, Qinviridae, Rhabdoviridae, Paramyxoviridae, Pneumoviridae, Lispiviridae, Bornaviridae, Nyamiviridae*, and *Filoviridae*. Most of these NSV families have non-segmented genomes. Exceptions are the *Chuviridae*, which have several different genome organizations, including non-segmented linear, non-segmented circular, bi-segmented linear, and bi-segmented circular. It is presently unclear if segmentation occurred relatively recently in this virus family and whether that may help explain their mixed genome organization and an RdRp-NP pair that is evolutionarily similar to the non-segmented *Haploviricotina*. In addition, we also observe that coffee ringspot virus (CRV), orchid fleck virus (OFV) and citrus chlorotic spot virus (CCSV), which are members of the bi-segmented *Dichoravirus* (a genus in the *Rhabdoviridae* family), cluster with the majority of the *Haploviricotina* in **Figure 6**.

The second main cluster indicated by the dendrogram consists of eight smaller clusters of NSV families with segmented genomes, at least in our dataset (see discussion for exceptions below). The first of these clusters (and second overall along the diagonal) groups three *Arenaviridae* family members and separates them from the other members of the *Bunyaviricetes* class. All ten *Articulavirales* order members cluster together in cluster 3. Similar to the NP MSA heatmap, the seven members of the *Naedrevirales* in our dataset form a single cluster (cluster 4) that is distinct from the large *Haploviricotina* subphylum cluster. Cluster 5 includes four members of the *Peribunyaviridea*, four of the *Fimoviridea*, as well as anopheles triannulatus orthophasmavirus (ATOPV), which is one of the four *Phasmaviridea* members in our dataset. The remaining clusters from left to right along the diagonal represent the four *Nairoviridea* (cluster 6), the *Hantaviridae* (cluster 7), the *Phenuiviridae* (cluster 8), and a final cluster consisting of the three remaining members of the *Phasmaviridae* (exclusive of ATOPV which is part of cluster 5) and the singular member of the *Tospoviridea* (cluster 9).

Overall, we find that the Alphafold2-derived data provide more resolution than the NP MSA analysis and that hierarchical clustering of the data produces NSV groups that are mostly consistent with the current *Negarnaviricota* phylogeny. Moreover, the NP AlphaFold2-based analysis is largely consistent with the RdRp MSA analysis, suggesting that the RdRp domain and NP sequences evolved together. Interestingly, and consistent with our RdRp and NP MSA analysis, most or all members of the *Naedrevirales* form a cluster that is distinct from the other *Haploviricotina*, the subphylum to which they have been assigned (37).

### PCA analysis of NP structures

To gain additional insight into the clustering of the *Naedrevirales*, we performed a principal component analysis (PCA) of our AlphaFold2 dataset. Using kmeans clustering (n=2) (**Figure 7**), we observe that the NPs encoded by viruses from the *Mononega*- and *Jingchuvirales* orders have a consistently high structural similarity with each other, while NPs encoded by the viruses of *Polyploviricotina* NSV subphylum and the *Naedrevirales* display greater structural variability and distinction between viral families, which is consistent with previous hypotheses (2). The seven *Naedrevirales* appear distinct from the other *Haploviricotina* as well as the *Polyploviricotina* NSVs. In accordance with the NP Alphafold2-based heatmap, the three members of *Dichoravirus* genus still cluster tightly within the *Haploviricotina* cluster, as does the segmented member of the *Chuviridea*. These PCA data together with the data presented in **Figure 5** and **6** suggest that the observed divergence of the *Naedrevirales* from the rest of the *Haploviricotina* is consistent and not solely attributable to the segmented nature of the genome.

**Figure 7:**
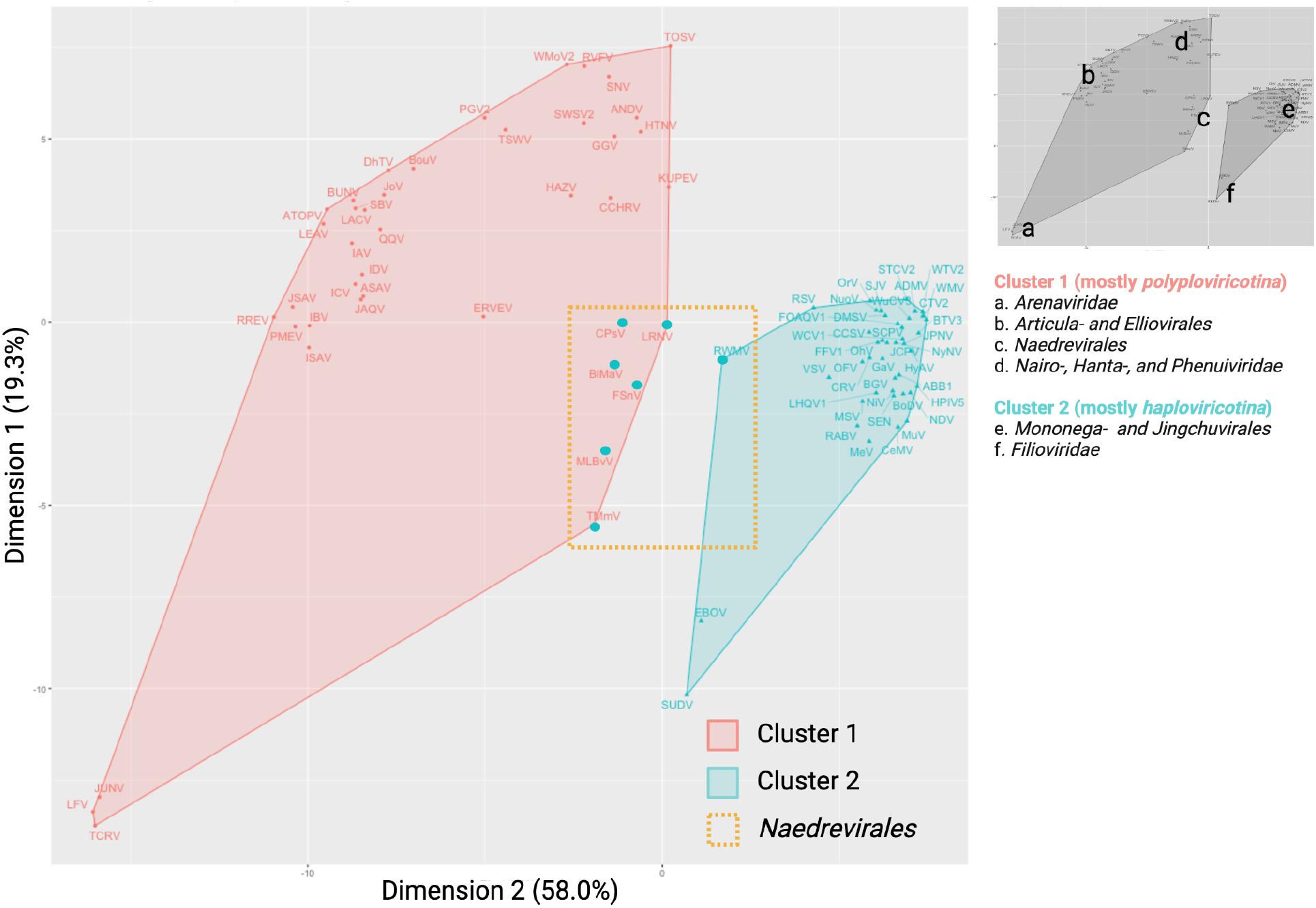
NP FATCAT kmeans clustering PCA plot. The NSV AlphaFold2-derived NP structural data was aligned using FATCAT, clustered by the kmeans method, and presented as a PCA plot. With the exception of six of the seven *Naedrevirales* (represented in the red *Polyploviricotina* cluster as large cyan data points), the clustering classifies each NSV according to the current ICTV subphylum: *Polyploviricotina* (red, cluster 1) and *Haploviricotina* (cyan, cluster 2). The *Naedrevirales* are highlighted with a yellow dashed-line box. The location of the various virus families and orders is indicated with numbers on the right.

## Discussion

Phylogenomic analyses of RNA virus sequences have been predominantly performed by comparing RdRp domain amino acid sequences, as this domain is universally conserved. A Clustal Omega-generated MSA of the NSV RdRp sequences shows a clustering that is consistent with the phylogenetic classification currently used by the ICTV at the family level (**Figure 1 and 5**) (27, 28). However, the *Naedreviridea* partly cluster separately from the *Haploviricotina*, the subphylum to which they have been assigned based on the predicted capping activity of the RNA polymerase. This may mean that the *Naedreviridae* are an outlier whose capping abilities evolved separately from the RdRp domain, or that they have been classified incorrectly based on limited molecular information. The RdRp MSA may also not be complete. Indeed, FSnV or TMmV are missing from the MSA because complete genome sequences, and in particular the RdRp domain sequences, are not available at this time. While we were not able to make a comprehensive assessment of all NSV incomplete genomes available, it is likely that other NSVs have been identified, but not yet or incorrectly classified due to the absence of RdRp or complete RNA polymerase sequences. We cannot rule out that the missing data may have affected the clustering of the *Naedreviridae* in our analysis.

Similar to the RdRp, the NP gene is present in all NSVs. When we generated an NP MSA for the same viruses as the RdRp MSA and included FSnV and TMmV, the percent identity matrix showed clustering of virus families consistent with the family clustering seen in the RdRp MSA. However, any virus order and class resolution was lost in the NP MSA (**Figure 5**). This observation is in agreement with other studies showing that the RdRp domain sequences provide the most evolutionary information for viral genome analyses. While sequence alignments help us to estimate the evolutionary relationships within NSVs, no previous work has been done to explore the potential of utilizing structural data to assess NSV phylogenetic patterns. It is well acknowledged that proteins are more conserved at the structural level than at the sequence level (11, 12). As structural data is becoming more readily available, the wealth of information it provides should be incorporated into our understanding of the functional conservation of viral proteins throughout time. Indeed, recent studies have shown that structure-based phylogenetic analyses can outperform sequence-based phylogenetic analyses (9–17).

We integrated AlphaFold2, AlphaPickle, FATCAT 2.0, and original data processing and screening, to build a pairwise structural alignment map of NSV NPs (**Figure 2**) (13, 31, 36). Even though we used the same amino acid sequences as for the MSA, the structural alignment allowed virus order and class clustering patterns to emerge. These clusters were largely identical to the RdRp domain MSA-based clusters (**Figure 6**). Only MLBvV clustered differently in the RdRp domain heatmaps compared to the NP heatmap, showing high similarity with the *Haploviricotina*, whereas the other *Naedrevirales* clustered with the *Polyploviricotina*. Interestingly, in our PCA analysis of the NP structure data (**Figure 7**), six of the seven *Naedrevirales* were assigned to the *Haploviricotina* cluster. Only the Ranunculus White Mottle Virus (RWMV) was assigned to the *Haploviricotina* cluster. Consequently, in all our analyses, the segmented *Naedrevirales* appear different from the *Haploviricotina*.

AlphaFold 2.0 utilizes available experimental data to make predictions about the structure of a given (unknown) protein sequence. Given that this deep learning method assumes the available data to be accurate, the results of the pairwise NP structure alignment may be biased towards current data and the currently accepted phylogeny that is based on these data. For instance, because there are structures available for *Rhabdoviridae* NPs, but no structures for *Naedrevirales* NPs, the predicted NP structures for the *Naedrevirales* may be inherently less accurate than NP structures for *Dichoravirus*, which appears more related to the *Rhabdoviridae*. To minimize this potential bias, only highly confident structures were utilized in this study. Moreover, we find that the observed *Naedrevirales* and *Chuviridae* clustering are supported by the RdRp MSA analysis, making it less likely that the observed clustering is an artifact of the lack of experimental structure data currently available. We suggest that the current ICTV classification of the *Naedrevirales* be critically examined in the future when more molecular biological data becomes available.

Electron microscopy analyses have shown that *Naedrevirales’* RNPs are flexible, having internally coiled loop-like structures, similar to *Elliovirales* RNPs, and more linear collapsed duplex structures, similar to *Articulavirales* RNPs (38). Morphologically, *Naedrevirales* RNPs most closely resemble RNPs of viruses in the *Tospoviridae* family (of the *Elliovirales* order), yet there is no evidence that the virions of *Naedrevirales* are enveloped. The same evidence is currently lacking for members of the *Tospoviridae* as well. Previous analysis of the overall architecture of NSV RNPs has shown a pattern in RNP flexibility that corresponds directly to genome organization (4). However, since there are only two micrographs available for *Naedrevirales* RNPs, it is possible that this observation is an outlier rather than consistent across all seven species. Obtaining additional electron micrographs is necessary to support a more holistic phylogenetic classification based on genome morphology, genome sequence, and/or protein structure analyses.

It has long been an outstanding question how genome segmentation evolved. The viral families that have been previously hypothesized to be associated with the evolution of NSV genome segmentation include *Naedrevirales* family along with viral species that belong to *Chuviridae* (6). While all currently identified species of *Naedrevirales* have segmented genomes, viral species with both non-segmented and segmented genomes have been identified within the *Chuviridae* family. Our current NP structure dataset includes seven non-segmented *Chuviridae* family members and one segmented *Chuviridae* family member. All eight have been assigned to the *Jingchuvirales* order. Consistent with this assignment, our analysis of the *Chuviridae* family members NPs, including the segmented WuCV3, shows a firm clustering with the non-segmented NSVs. Consequently, RdRp and NP analyses yield similar evolutionary relationships and it is tempting to conclude that there is no clear predictor of segmentation based on either analysis of the RdRp sequence or NP structure.

As has been proposed elsewhere, it is possible that segmented NSV genomes evolved multiple times. On the one hand, this means that segmentation could have evolved independently in multiple virus lineages. On the other hand, it means that once segmentation evolved once in one lineagues, further segmentations could evolve. The latter hypothesis could explain the large differences seen between *Articulavirales* and *Elliovirales* in terms of the number of genome segments and the gene organization in the respective viral genomes. This former hypothesis would be consistent with the *Naedrevirales* and *Chuviridae* data. Moreover, it has been observed that several viral species that belong to the *Monogenavirales* order have segmented genomes, including orchid fleck dichorhavirus (OFV), which is classified within the *Rhabdoviridae* family and possesses two genome segments (6). Phylogenetic analysis suggests that this virus evolved from a non-segmented plant virus within the *Rhabdoviridae* family (39). Our data shows that OFV NP, as well as CFV and CCSV NPs, are homologous to the NPs from other non-segmented *Rhabdoviridae* members, despite the segmented character of their genome. Interestingly, the non-segmented *Tulasviridae* were recently identified as members of the *Elliovirales*. In our analysis, the *Tulasviridae* NP did not meet the quality cut-offs for analysis (see **Supplemental Information**).

While we can speculate about the origin of segmentation, recent large-scale virus discoveries have demonstrated that we always need to consider the likelihood that viral species remain to be identified, and that the “origin species” of NSV genome segmentation has not been discovered yet. It is also possible that this species has been identified, but that it has so far been excluded from large scale evolutionary analyses because genome sequences were incomplete or missing. In particular for the latter aspect, the findings presented here suggest a useful role for including structural information based on sequences of non-RdRp proteins in phylogenomic analyses. We therefore hope that our findings will help inspire new research and ultimately a better understanding of NSV evolution and a more intuitive, consistent classification.

## Materials and Methods

### Data Acquisition

Sequences and currently accepted taxonomic classifications were acquired from ICTV and NCBI sequence databases. Accession numbers are available in **Supplementary Table 1**.

### Clustal Omega MSA

The EMBL-EBI Clustal Omega Multiple Sequence Alignment web server was used to perform MSA analysis for both the RdRp and the NP alignments (40). Protein sequences were consolidated into a single FASTA file and run with ClustalW output parameters and restricted to the predefined order of input. The percent identity matrix was used to make the MSA heatmaps in R.

### AlphaFold 2.0 structure prediction

The newest AlphaFold algorithm (version 2.0) was used on the Della HPC at Princeton University to predict the structures of all NPs within the dataset (13). The general performance of AlphaFold2 with viral NPs was assessed using the 35 known NP structures deposited in the PDB. The amino acid sequences of the viral NPs were used as the inputs and a maximum PDB template date was defined to be before the release date of each solved structure to ensure AlphaFold2 could not “cheat” during prediction.

The predicted NP models were then assessed for confidence (with both pLDDT and PAE metrics) using AlphaPickle (36). The predicted structures were then compared to the experimentally solved structures deposited in the PDB and all predicted NP structures were determined to be statistically accurate. The following structural analyses with the expanded dataset (including 84 NSVs) used all AlphaFold2 predicted NP structures to reduce structural solution method bias.

### Intrinsic protein structure disorder prediction

The IUPred package was downloaded from https://iupred.elte.hu/ to allow for faster data acquisition (26). Single sequences can be analyzed through the freely accessible web-server.

### FATCAT 2.0 Structure Alignment

The source code for Flexible structure AlignmenT by Chaining Aligned fragment pairs allowing Twists (FATCAT) was retrieved from the GodzikLab github repository (31). The software was compiled and built on the Della HPC at Princeton University. Original data organization code produced an input list of NP pairs in the predefined order. The FATCAT software then used the AlphaFold predicted NP structures to run flexible structure alignment for the 3,486 defined unique pairs. The results were quantified as a p-value, or the probability of observing a more identical alignment between the structure pairs. The alignment data was illustrated as a heatmap to facilitate clustering comparison to the MSA percent identity heatmaps.

### Alignment analysis in R

Original code to process and visually analyze the resulting alignment data was written in R. Heatmaps were created using R package ggplots.

### Clustering and PCA analysis in R

The FATCAT structure alignment data was clustered by dissimilarity using hierarchical clustering with the Ward agglomeration method from the stats R package. Principal component analysis (PCA) of the FATCAT structure alignment data was performed using the kmeans clustering algorithm with n=2 clusters in the stats R package.

## Data Availability and Requirements

Predicted AlphaFold structure files are available in pdb format in the supplementary information. Dataset acquisition codes can be found in **Supplementary Table 1**.

## Supplementary Information

Supplementary Table 1: Full NSV Dataset

Supplementary File 1: Initial Dataset NP PDBs (.pdb files)

Supplementary File 2: Full NSV Dataset NP AlphaFold PDBs (.pdb files)

Supplementary File 3: Raw data files

## Raw data files

1. AlphaFold2 performance alignment data: AlphaFold_Performance_NSV_NPs
2. IUPred2 NP gene raw data: IUPred2_NPgene_Rawdata.xlsx
3. RdRp MSA percent identity matrix raw data: RDRP_MSA_PIM.xlsx
4. NP MSA percent identity matrix raw data: NP_MSA_PIM.xlsx
5. FATCAT alignment raw data: NSV84_FATCAT.aln

## Acknowledgements

The authors would like to thank Dr. Ned Wingreen, Dr. Maria Rosenthal and Dr. Ingrida Olendraite for insightful discussions regarding the evolution of NSVs, as well as members of the Wingreen lab for discussion of the computational analyses. KRS was supported by NIH grants R01 GM140032 (to Ned Wingreen) and R01 AI170520 (to Adam Lauring, AJ te Velthuis and Colin Russell). ATV was supported by NIH grants DP2 AI175474 (to AJ te Velthuis) and R01 AI170520 (to Adam Lauring, AJ te Velthuis and Colin Russell), and Wellcome Trust and Royal Society grant 206579/17/Z (to AJ te Velthuis).

The authors are pleased to acknowledge that the work reported on in this paper was performed using the Princeton Research Computing resources at Princeton University which is a consortium of groups led by the Princeton Institute for Computational Science and Engineering (PICSciE) and Office of Information Technology’s Research Computing.

