## Supplemental File 2 for "Using structure prediction of negative sense RNA virus nucleoproteins to assess evolutionary relationships": draft_Proof_hi.pdf



| | |
|---|---|
| Journal: | *Virus Evolution* |
| Manuscript ID | Draft |
| Manuscript Type: | Research Article |
| Date Submitted by the Author: | n/a |



| | |
|---|---|
| Complete List of Authors: | Sabsay, Kimberly; Princeton University, Department of Molecular Biology<br>te Velthuis, Aartjan; Princeton University, Department of Molecular Biology |



| | |
|---|---|
| Keywords: | negative-sense RNA virus, nucleoprotein, AlphaFold 2.0, RNA polymerase, evolution, segmentation |


Note: The following files were submitted by the author for peer review, but cannot be converted to PDF. You must view these files (e.g. movies) online.

Supplementary_File_1_Initial_Dataset_NP_PDBs.zip
Supplementary_File_2_Full_NSV_Dataset_NP _AlphaFold_PDBs.zip

SCHOLARONE™ Manuscripts

### Using structure prediction of negative sense RNA virus nucleoproteins to assess evolutionary relationships

Kimberly R. Sabsay[1,2] and Aartjan J.W. te Velthuis[1,*]

[1] Lewis Thomas Laboratory, Department of Molecular Biology, Princeton University, Princeton, NJ 08544, United States.
[2] Sigler Institute, Princeton University, Princeton, NJ 08544, United States.

For Review Only

**Key words:** negative-sense RNA virus, NSV, nucleoprotein, NP, RNA polymerase, RdRp, phylogenomics, evolution, segmentation, genome, AlphaFold 2.0

#### Abstract

Negative sense RNA viruses (NSV) include some of the most detrimental human pathogens, including the influenza, Ebola and measles viruses. NSV genomes consist of one or multiple single-stranded RNA molecules that are encapsidated into one or more ribonucleoprotein (RNP) complexes. Current evolutionary relationships within the NSV phylum are based on alignment of conserved RNA-dependent RNA polymerase (RdRp) domain amino acid sequences. However, the RdRp-based phylogeny does not address whether other core proteins in the NSV genome evolved along the same trajectory. Moreover, the current classification of NSVs does not consistently match the segmented and non-segmented nature of negative-sense virus genomes. Viruses belonging to e.g. the *Serpentovirales* have a segmented genome but are classified among the non-segmented negative-sense RNA viruses. We hypothesized that RNA genome segmentation is not coupled to the RdRp domain, but rather to the nucleocapsid protein (NP) that forms RNP complexes with the viral RNA. Because NP sequences are too short to infer robust phylogenetic relationships, we here used experimentally-obtained and AlphaFold 2.0-predicted NP structures to probe whether evolutionary relationships can be estimated using NSV NP sequences and potentially improve our understanding of the relationships between NSV subphyla and the NSV genome organization. Following flexible structure alignments of modeled structures, we find that the structural homology of the NSV NPs reveals phylogenetic clusters that are consistent with the currently accepted NSV taxonomy based on RdRp sequences with one key difference: the NPs of the segmented *Serpentovirales* cluster with the other segmented NSV. In addition, we were able to assign viruses for which RdRp sequences are currently missing to clusters of structural similarity. Overall, our results suggest that the NSV RdRp and NP genes largely evolved along similar trajectories, that NP-based clustering is better correlated with the NSV genome structure organization, and that even short pieces of genetic, protein-coding information can be used to generate pair-wise comparisons, potentially making metagenomic analyses more valuable.


#### Introduction

Negative sense RNA viruses (NSVs) are important human pathogens, and include the influenza A virus (IAV), Rift Valley fever virus (RVFV), Lassa virus (LASV), measles virus (MeV), rabies virus (RABV), and Ebola virus (EBOV). The genomes of NSVs consist of single-stranded, negative sense RNA that is copied in the context of a ribonucleoprotein (RNP) complex. Each RNP is composed of a viral RNA (vRNA) template, an RNA polymerase, and numerous nucleoproteins or nucleocapsids (NPs) (1, 2). The self-oligomerizing nucleoprotein (NP) molecules make up the majority of each RNP. These NPs protect the genome from degradation, act as scaffold for RNA structure formation, and assist in viral replication by acting as processivity factor (3, 4). Despite these conserved features, both the genome organization, and replication and transcription mechanisms vary widely among NSVs (5). Understanding the properties and evolutionary history of NPs may shed light on how different NP support NSV genome structures and RNA polymerase processivity.

The International Committee on Taxonomy of Viruses (ICTV) established the phylum *Negarnaviricota* for negative sense RNA viruses (NSVs) in 2019 (**Figure 1)**. Every virus within

*Negarnaviricota* shares a common three-gene core: the RNA polymerase gene(s), membrane glycoprotein genes, and an NP gene (6). The RNA-dependent RNA polymerase (RdRp) domain of the RNA polymerase gene product is approximately 300 amino acids long and conserved amongst all RNA viruses (kingdom *Orthonavirae*), whereas the rest of the RNA polymerase is not. These characteristics have made the RdRp domain the focal point for evolutionary analyses (7, 8). The RdRp domain phylogenies have mapped out five major branches of *Orthonavirae* and elucidated ancestral trends amongst them. While traditional phylogenomics utilizes metagenomics data, sequence alignments, and comparison metrics, the ever-expanding wealth of structural information provides an avenue for the refinement of important branching points (9–17). In particular, structural comparisons of RdRp domains have illustrated surprising similarity amongst *Orthomyxoviruses* (with negative-sense ssRNA), flaviviruses (with positive-sense ssRNA) and *Cystoviruses* (with dsRNA), suggesting NSVs originated from dsRNA viruses which in turn evolved from positive-sense ssRNA viruses (14). NSVs subsequently evolved into sub-phyla (**Figure 1**), although the details of this split are unknown.

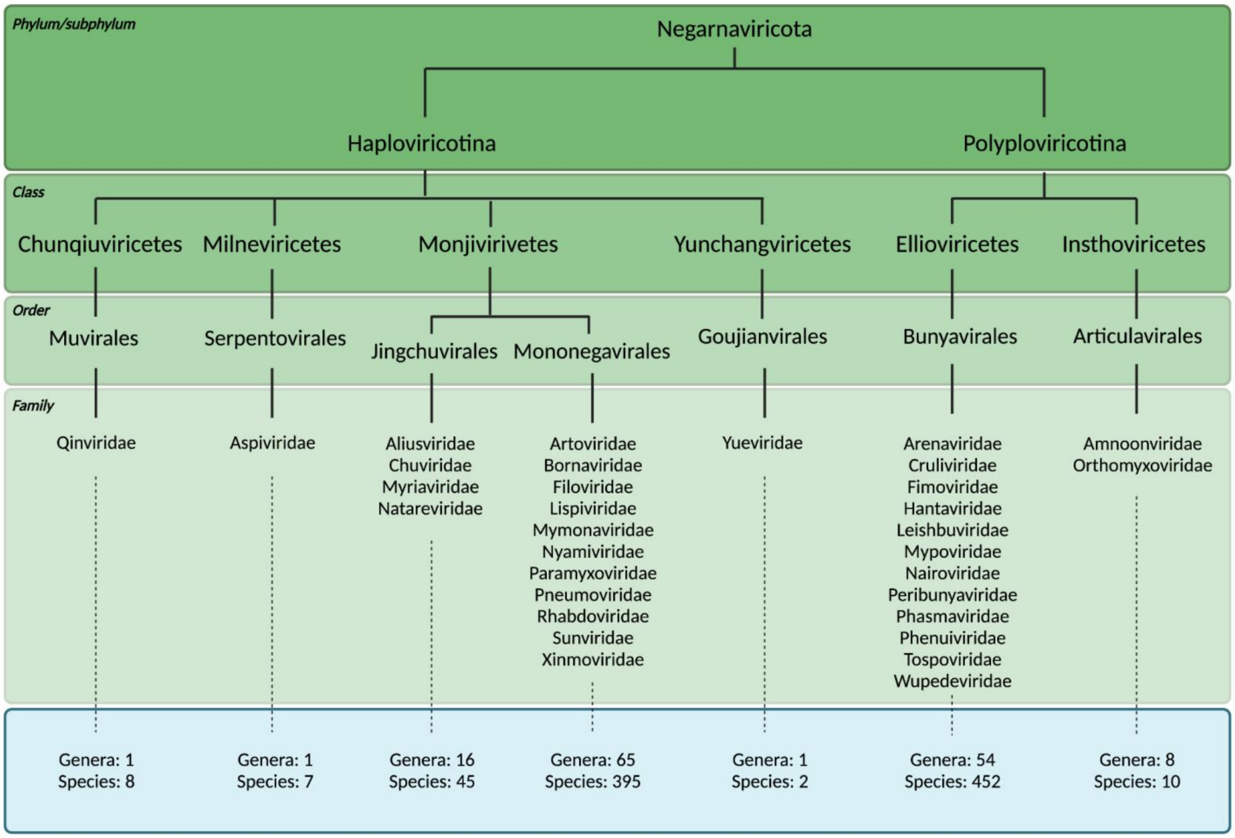


**Figure 1: Classified viral species within *Negarnaviricota*.** Taxonomic breakdown of the *Negarnaviricota* phylum of the *Riboviria* realm according to the ICTV and NCBI databases. The phylum consists of all NSVs and is divided into the subphyla *Haploviricotina* and *Polyploviricotina* based on the presence or absence of mRNA capping activity in the RNA polymerase and the organization (non-segmented or segmented) of the viral genome. Note that the *Serpentovirales* have a segmented genome, but an RNA polymerase with mRNA capping activity.

NSVs have historically been connected evolutionarily through the conserved RdRp domain (18). Based on the RdRp domain sequence, the NSV phylum is subdivided into *Haploviricotina* and

*Polyploviricotina*. This division separates the NSVs that have a non-segmented genome and an RNA polymerase that possess mRNA capping activity from those that have a segmented genome and an RNA polymerase that performs cap-snatching to cap viral mRNAs, respectively (8). Genome segmentation allows for reassortment of gene segments, which can contribute to global pandemics and the spread of immune or antiviral resistance mutations (19, 20). Segmentation also makes the differential expression of viral genes easier to regulate (21, 22). On the other hand, segmentation complicates virus genome packaging. A virion of a segmented virus is not viable unless it contains the entire set of viral genome segments. The coordination and packaging of all segments is thus a crucial step in segmented NSV infections. In influenza A viruses, the majority of virion particles contain one copy of each of the eight genome segments (23).

Previous work has used analysis of RdRp domain sequences to understand the evolution of genome structure and segmentation within *Negarnaviricota* or the relations among NSV orders (6, 24). The important role of NPs in vRNA protection, replication, and packaging, as well as in RNP morphology, suggests that the evolutionary history of NPs may reveal additional insights into the evolution of NSV genome segmentation. However, NP sequences are too short for robust phylogenetic analyses. We here combined experimentally obtained structural data and the deep learning tool AlphaFold 2.0 (AlphaFold2) (**Figure 2**) to explore if the evolutionary relationships among NSVs can be reconstructed from NP sequences and if this relationship provides additional insight into the divergence of the NSV subphyla and the emergence NSV genome segmentation. In addition, this approach may be useful for metagenomics and virus discovery studies where RdRp sequences may not be complete or missing. We find that the structural homology of NPs within NSVs provides more information for pair-wise similarity comparisons than NP sequences alone. In addition, we observe that the NP structural clustering largely matches the clustering and known inferred phylogeny of the RdRp domain, with the exception of the segmented *Serpentovirales*, suggesting that the two NSV core genes largely co-evolved. The clustering of the *Serpentovirales* with the other segmented NSVs, instead of the non-segmented NSVs, suggest that the NP structure-based clustering is better correlated with the division between the segmented and non-segmented NSV genome organization than the RdRp-based phylogeny.

For Review Only

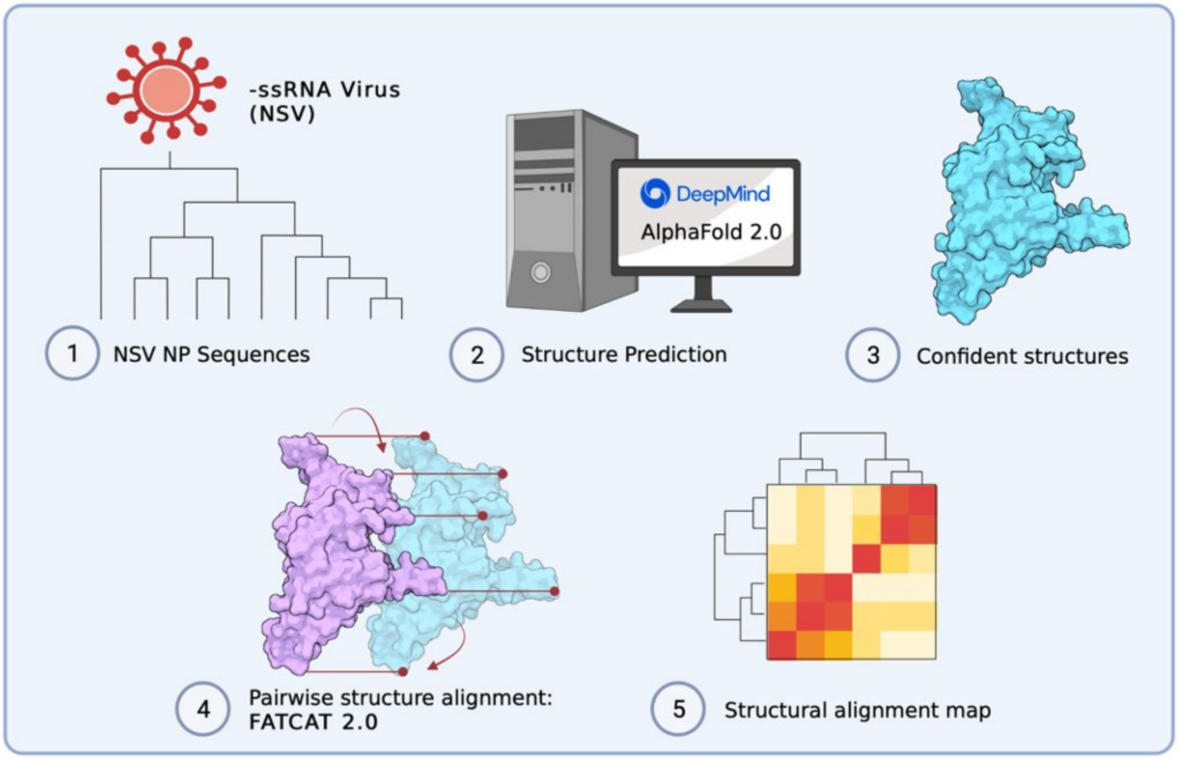


**Figure 2: Computational pipeline to build structural alignment map.** (**1-2**) Data was acquired based on sequence availability. AlphaFold 2.0 was used to predict structures. (**3-4**) AlphaPickle was used to visualize confident metrics, following which structures were selected based on confidence score cut-offs and individual pairwise flexible alignments were computed using the FATCAT 2.0 package compiled on a high performance cluster (HPC). (**5**) Data processing was performed to parse structural alignment data into a concise heatmap for comparison with MSA percent identity matrices.

Review Only

#### Results

##### *Experimental NP structure dataset*

At the time of submission of this manuscript, there were 457 viral species in the *Haploviricotina* and 462 species in the *Polyploviricotina* assignments of the ICTV database. The taxonomical breakdown is summarized in **Figure 1**. Structural data for NPs was found for 34 of these NSVs in the PDB database. Within this initial dataset, 21 NP structures were from *Polyploviricotina* and 13 from *Haploviricotina*. Given the relatively even ratio of officially classified viral species in each subphyla in the ICTV, this initial NP structure dataset appears to be biased towards the human-disease causing, segmented NSVs (**Figure 3**).

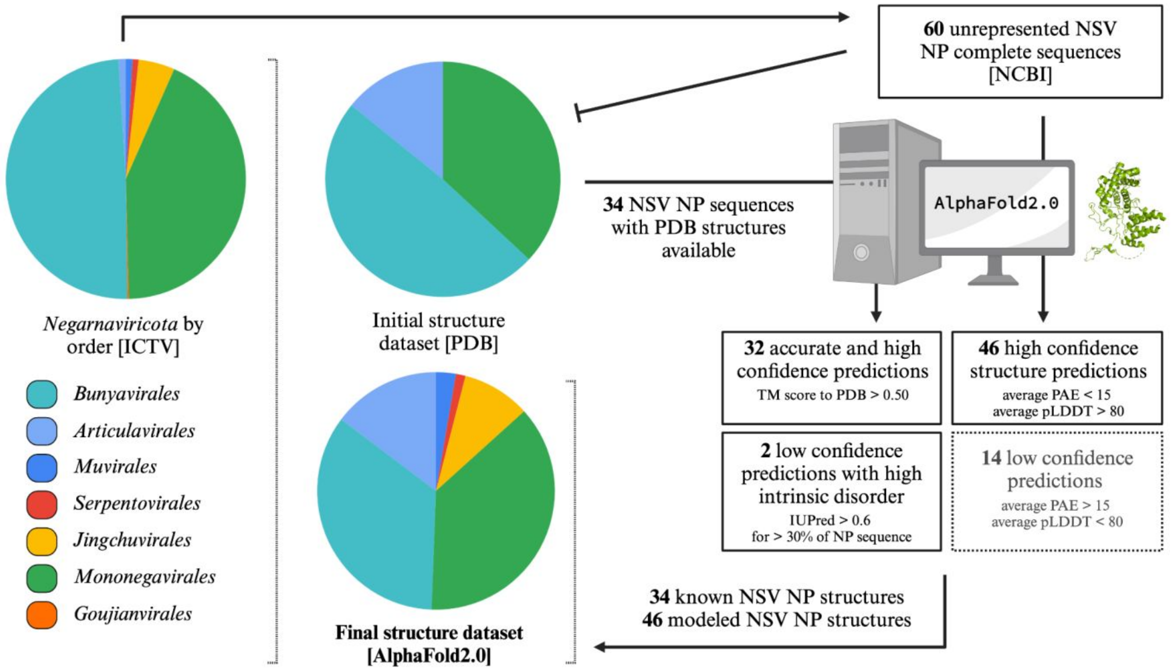


**Figure 3: Flowchart for the creation of a representative structural dataset of Negarnaviricota NPs.** The composition breakdown of *Negarnaviricota* by phylogenetic order (*left*) was determined by the number of species in each order in the ICTV/NCBI database at the time of the manuscript preparation. The percentages of each order are as follows: 49.2% *Bunyavirales*, 1.0% *Articulavirales*, 0.9% *Muvirales*, 0.8% *Serpentovirales*, 4.9% *Jinchuvirales*, 43.0% *Mononegavirales*, and 0.2% *Goujianvirales*. At the time of the manuscript preparation, there were 34 NSV NP structures available in the PDB database with representation from only the *Bunyavirales*, *Articulavirales,* and *Mononegavirales*. The final NSV NP dataset included a total of 80 structures generated with AlphaFold 2.0.

For Review Only

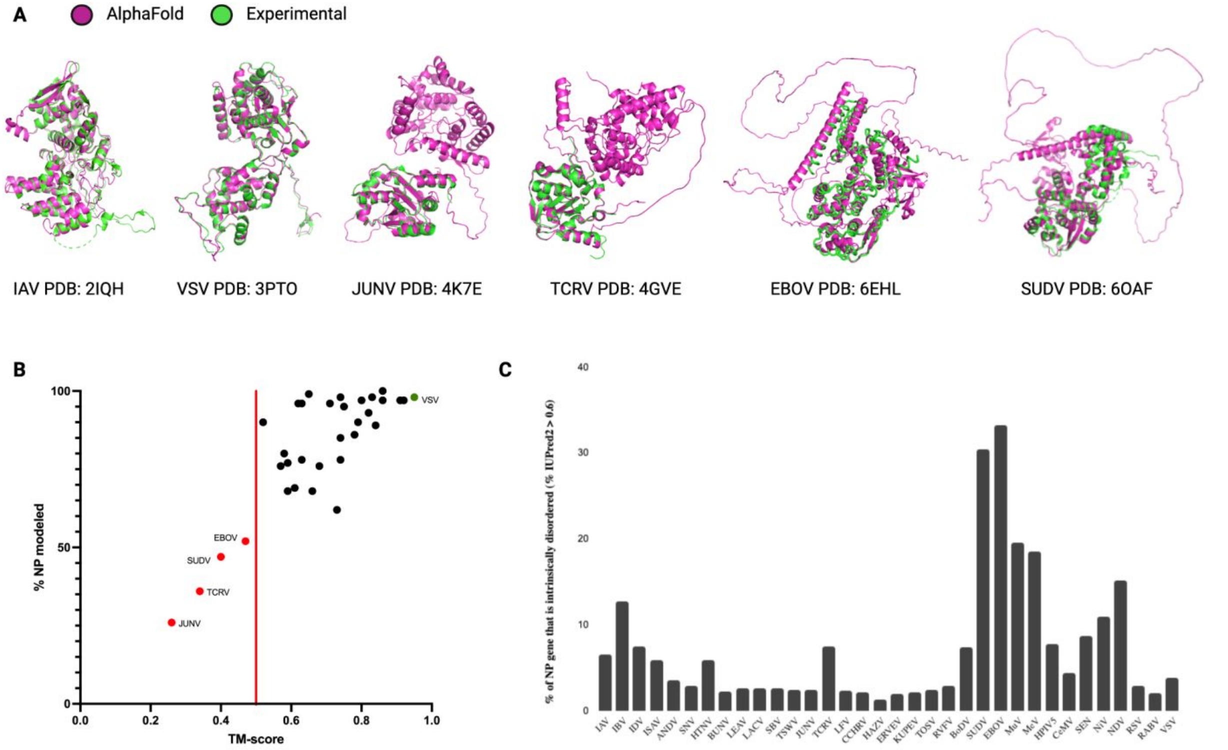


**Figure 4: AlphaFold2 structure prediction performance on solved NSV nucleoproteins.** Overall prediction performance of AlphaFold 2.0 on NSV NPs was assessed using an initial dataset of 34 NSVs with experimentally solved structures deposited in PDB. (**A**) Structural alignments of AlphaFold2 predictions (magenta) and experimental structures (green) are displayed for the most accurate prediction (VSV and IAV, left), a partially resolved experimental structure with the lowest % of NP modeled (JUNV and TCRV, middle), and a highly intrinsically disordered structure (SUDV and EBOV, right). (**B**) The predicted AlphaFold 2.0 structures were compared to the experimental NP structures using rigid jFATCAT structural alignment and plotted according to the alignment score (TM-score). Four structures fell below a TM-score of 0.50 (red line) and are depicted as red points and labeled. The top scoring AlphaFold prediction with an almost perfect alignment (VSV NP) is denoted by the green data point. (**C**) Intrinsic protein disorder predictions for the initial dataset using IUPred2 show a high percentage of disorder within the EBOV and SUDV NPs.

##### *Expanding the NP structure dataset using AlphaFold2*

We next used AlphaFold2 to extend the experimental NP structure dataset to obtain better structural coverage of the *Negarnaviricota* NPs. To this end, additional viral species with known and complete NP sequences were chosen from unrepresented NSV families and the NP structures of these viruses predicted using AlphaFold2. The structure predictions that passed the confidence requirements (average PAE scores <15 and average pLDDT scores >80) were included in the final NP structure dataset (**Figure 3**). This dataset contained a total of 80 NP structures with 50.6% from *Haploviricotina* and 49.4% from *Polyploviricotina* (**Supplementary Table 1**).

##### *Analysis of the NP and RdRp domain MSAs*

To assess if the NP structures contained enough information to investigate the evolutionary history of NP, we first generated an MSA of the RdRp domain sequences to use as a benchmark. Only 78 of the 80 NSVs were included in this MSA as the freesia sneak virus (FSnV) and tulip mild mottle virus (TMmV) RdRp sequences were not available in sequence databases at the time of analysis. The MSA rows were arranged according to the current ICTV classification and the resulting MSA used to generate a percent identity matrix. This matrix is shown as a heatmap in **Figure 5A, left**. We next generated an MSA using the NP sequences and illustrated the NP percent identity matrix as a heatmap as well (**Figure 5A, right**).

For Review Only

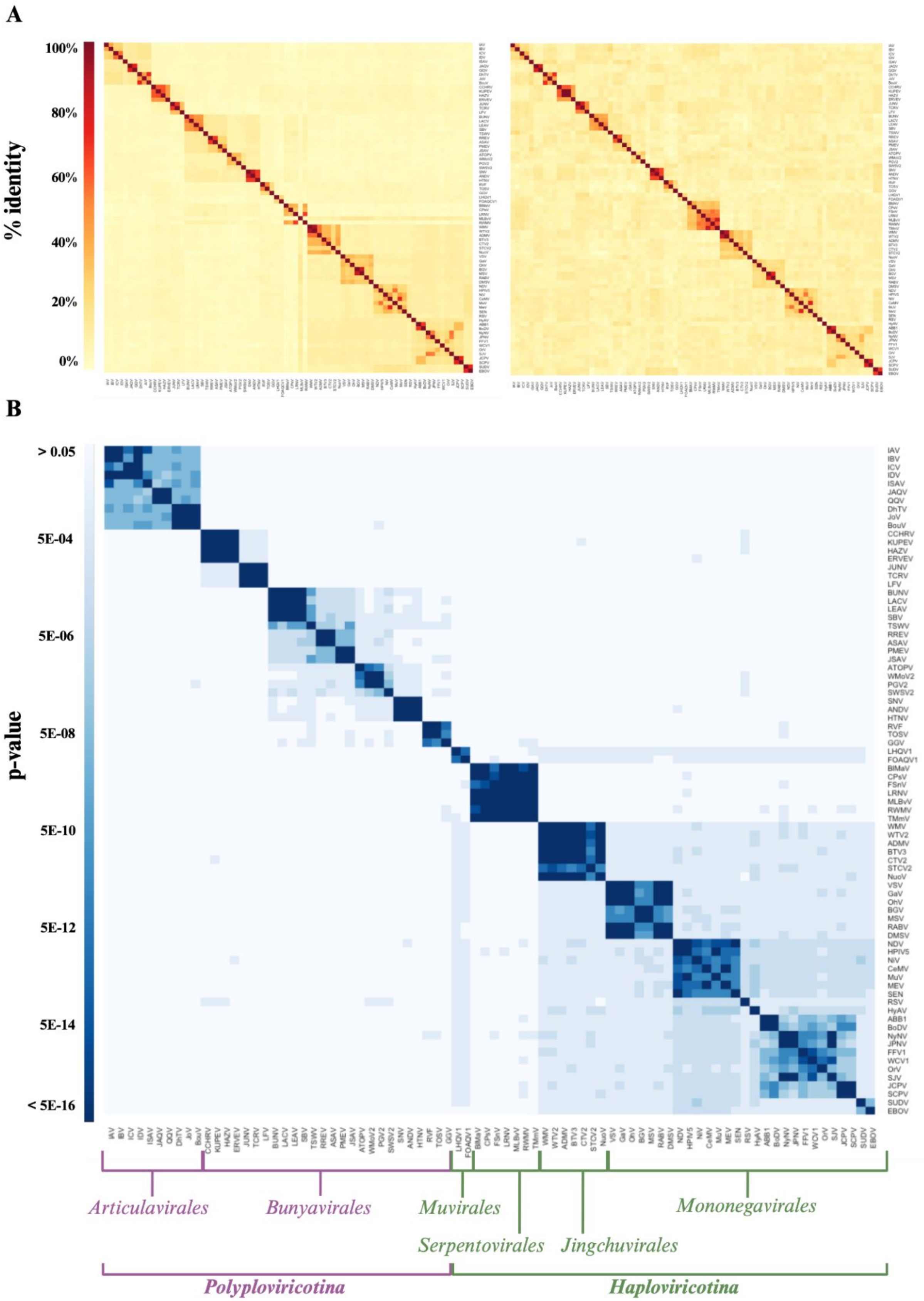
A
% identity
100%
80%
60%
40%
20%
0%
B
p-value
> 0.05
5E-04
5E-06
5E-08
5E-10
5E-12
5E-14
< 5E-16
IAV IBV ICV IDV ISAV JAQV QQV DhTV JoV BouV CCHRV KUPEV HAZV ERVEV JUNV TCRV LFV BUNV LACV LEAV SBV TSWV RREV ASAV PMEV JSAV ATOPV WMoV2 PGV2 SWSV2 SNV ANDV HTNV RVF TOSV GGV LHQV1 FOAQV1 BiMaV CPsV FSnV LRNV MLBvV RWMV TMmV WMV WTV2 ADMV BTV3 CTV2 STCV2 NuoV VSV GaV OrV BGV MSV RABV DMSV NDV HPIV5 NiV CeMV MuV MEV SEN RSV HyAV ABB1 BoDV NyNV JPNV FFV1 WCV1 OrV SJV JCPV SCPV SUDV EBOV
Articulavirales
Bunyavirales
Muvirales
Mononegavirales
Serpentovirales
Jingchuvirales
Polyploviricotina
Haploviricotina

**Figure 5: NSV NP structural alignment matrix matches RdRp MSA matrix**. (**A**) RdRp (*left*) and NP (*right*) amino acid sequence MSAs illustrated as percent identity matrices. Note that the RdRp MSA did not include 2 of the 80 NSVs used in other matrices, as freesia sneak virus (FSnV) and tulip mild mottle virus (TMmV) RdRp sequences were available in sequence databases at the time of analysis. (**B**) NP structural alignment by FATCAT in an order consistent with the heatmaps in panel A, and illustrated as a p-value matrix.

The RdRp and NP heatmaps shown in **Figure 5A** are presented with an identical x- and y-axis order, with the exception of FSnV or TMmV, which are absent in the RdRp MSA heatmap. The clustering can thus be qualitatively compared across the two alignments. However, it is important to note that the color scales of the MSA and FATCAT heatmaps should not be directly compared as they derive from different alignment calculations. The predefined order of the heatmaps separates the segmented NSVs (first 36, starting left) from the non-segmented NSVs (last 44). The heatmap that is based on the RdRp MSA shows four dark clusters in the top left corner that are grouped into a single larger cluster. This larger group consists of 10 NSVs that all belong to the *Articulavirus* order. The four smaller clusters represent the *Influenza virus* (4), *Quaranjavirus* (2), *Isavirus* (1), and *Thogotovirus* (3) genera, in line with the current ICTV phylogeny. Comparing the RdRp and NP MSA heatmaps, we note that while the four virus family clusters are visible in the NP heatmap, the virus order clustering seen in the RdRp heatmap is lost in the NP heatmap.

In the RdRp heatmap, the next two darker clusters correspond to the four *nairoviruses* and three *arenaviruses*, which all belong to the *bunyavirus* order. The NP heatmap shows the same *nairovirus* and *arenavirus* clustering, but neither heatmap clusters the two families into the same order or even a larger cluster. The next clusters in the RdRp heatmap are made up of 16 additional *bunyaviruses* that fall into the five viral families *Peribunyaviridae* (4), *Tospoviridae* (1), *Fimoviridae* (4), *Phasmaviridae* (4), and *Hantaviridae* (3). The RdRp and NP heatmaps cluster these 16 viruses into the same five families as the current taxonomy, but only the RdRp heatmap clusters them into a single larger cluster. The three remaining bunyaviruses in our data set belong to the *Phenuiviruses* family. We find them clustered adjacent to the above larger cluster of five bunyavirus families. We do not observe a single cluster that perfectly encapsulates all members of the *Bunyavirus* order.

From GGV onwards, we find a small cluster of two *Muvirales* and a larger cluster of *Serpentoviruses* in the RdRp (5) and NP heatmaps (7). The RdRp heatmaps clusters 4 of the 5 *Serpentoviruses* with the larger cluster of six *Bunyaviruses* families (the *Nairovirus* and *Arenavirus* are not included in this cluster). The exception is *Mirafiori lettuce big vein virus* (MLBvV), which the RdRp heatmaps clusters with a large *Haploviricotina* (non-segmented NSVs) cluster. In the NP heatmap, this clustering is not observed. The NP heatmap does show clustering of FSnV or TMmV, for which RdRp domain sequences were not available, with the *Serpentoviruses*. It is important to note here that, at the time of analysis, the *Serpentoviruses* are taxonomically classified within *Haploviricotina* based on the mRNA capping ability of the RNA polymerase even though their genomes are segmented. The RdRp domain MSA suggests that in spite of the capping domain, their NTP incorporation function may be more closely related to the *Bunyavirales*.

For Review Only

The second part of the heatmaps consists of 35 non-segmented NSVs that together form the clusters: *Rhabdoviruses* (7), *Chuviruses* (7), *Paramyxoviruses* / *Pneumovirus* (8), Qinvirus / Lispivirus (3), *Nyamiviruses* (6), *Bornaviruses* (2), and *Filoviruses* (2). In the RdRp domain heatmap, these seven clusters are combined into one large cluster, but this clustering is not evident in the NP heatmap. Overall, this analysis suggests that the virus family clusters are resolved in both the RdRp and NP heatmap and that the family clustering is consistent with the current phylogeny of *Negarnaviricota* (27, 28). However, at a higher level, the RdRp domain heatmap is able to identify some orders that the NP heatmap cannot. Thus, the RdRp domain sequence appears to offer more information to perform pair-wise similarity comparisons than the NP sequence.

***Pair-wise analysis of NP structures is comparable to RdRp domain MSA***

We hypothesized that the NP AlphaFold2 structure dataset contained more phylogenetic information than the NP sequences alone, because protein structures are generally more conserved than the primary sequence. To explore this, we had to align our AlphaFold2 NP structures. Aligning structural information requires a completely different approach and significantly more computational power than aligning primary amino acid sequence data. Foundational structural alignments utilize rigid body backbone algorithms. While this is a powerful approach for assessing the similarity of closely related structures that are solved in similar conditions, the natural motion and dynamics of protein structures are not accounted for in rigid body alignments (29, 30). As proteins can adopt multiple conformations according to their function or protein structures have different energy minima, it is critical to analyze structures in a way that tests whether two proteins identical, even if they have adopted different conformations (e.g. “open” or “closed”, or “active” or “inactive”). Various complex algorithms have been developed to attempt to incorporate dynamic motion into structural alignments. For our NP AlphaFold2 structure analyses, we used the Flexible structure AlignmenT by Chaining Aligned fragment pairs allowing Twists (FATCAT) algorithm (31). This method utilizes fragment pairs and iterative rotations/twists to converge on an alignment that would account for alternate conformations. As is the case with NSV NPs, the two-domain NP structures have been proposed to twist upon oligomerization and/or RNA-binding to form the helical chains seen in RNP complexes (32–35).

Pairwise alignments for all possible pairs of NPs within the dataset were individually computed. Each alignment pair resulted in an alignment p-value, which we interpret as the probability of observing a more identical alignment between the structure pairs. A structural alignment heatmap was generated from the alignment p-values and organized in the same predefined order as illustrated in the MSA heatmaps for direct comparison (**Figure 5C**). The diagonal represents self-alignment, or a p-value of exactly zero. A computational pipeline integrates AlphaFold2, AlphaPickle, FATCAT 2.0, and original data processing and screening code, to allow for high-throughput expansion of the pairwise structural alignment map described above (**Figure 2**) (13, 31, 36).

The first main cluster of the NP structure alignment agrees with the RdRp MSA, containing the 10 *articulaviruses*/*orthomyxoviridae*. The order/family cluster can be divided into four sub-clusters that match the four genera in our dataset. The following two main structure alignment clusters are

For Review Only

also consistent with the RdRp MSA and correspond to the *nairovirus* and *arenavirus* clusters. The remaining 19 bunyaviruses form one larger cluster that contains subclusters that match the six bunyavirus families. The NP structural alignment thus outperforms the NP MSA and generates a larger bunyavirus cluster that is not observed in the RdRp MSA.

The *Muvirales* follow the *Bunyavirales* in the NP structure alignment heatmap. Interestingly, the two included NPs appear to cluster with the large *Haploviricotina* cluster. The large cluster that follows the Muvirales in the NP structure alignment corresponds to the *Serpentoviruses*. In the RdRp MSA, MLBvV seemed to deviate from the other *Serpentoviruses* and cluster with the non-segmented viruses, whereas the other *Serpentoviruses* appeared to cluster with the *bunyaviruses*. In the NP structure alignment, MLBvV does not appear to be an outlier, but clusters tightly with the other *Serpentoviruses*, and remains separate from the non-segmented NSV cluster. The final main cluster within the NP structure alignment corresponds to the remaining non-segmented NSVs. This observation is fascinating as it contradicts the current classification of the *Serpentoviruses* as a member of *Haploviricotina* based on RdRp function and phylogenetic analyses and suggests for these viruses RdRp and NP functions diverged, which aligns with the capping capabilities of the RNA polymerase in combination with the segmented genome (37).

Overall, NPs encoded by viruses in the non-segmented NSV subphylum have a consistently high structural similarity with each other, while NPs encoded by viruses of the segmented NSV subphylum display greater structural variability and distinction between viral families, which is consistent with previous hypotheses (2). The fact that *Serpentoviruses* appear to encode NPs with structures that are structurally more consistent with NPs encoded by viruses from the segmented subphylum is a new insight and contradicts analyses performed using RdRp sequences. The currently accepted classification of NSVs fails to accurately differentiate NSV based on the structural organization of their genetic material. Our new analysis based on NP structural modeling reflects clustering consistent with the NSV genome organization.

#### Discussion

Phylogenomic analyses of RNA virus sequences are typically performed by comparing RdRp domain amino acid sequences. A Clustal Omega-generated MSA of the NSV RdRp sequences shows a clustering that is consistent at the family level with the phylogenetic classification currently used by the ICTV (**Figure 5A**) (27, 28). However, the RdRp MSA does not include all possible NSV sequences available. For instance, FSnV or TMmV are missing because complete genome sequences, and in particular the RdRp domain sequences, are not available at this time. When an NP MSA was performed for the same viruses as the RdRp MSA, the percent identity matrix showed some clustering of closely related NSVs, consistent with clustering seen in the RdRp MSA. However, any resolution of the virus order and class was lost in the NP MSA (**Figure 5A**). This observation agrees with other studies showing that the RdRp domain sequences provide the most evolutionary information for viral genome analyses. While sequence alignment helps us estimate the evolutionary relationships within NSVs, no previous work has been done to explore the potential of utilizing structural data to assess NSV phylogenetic patterns. It is well acknowledged that proteins are more conserved at the structural level than at the sequence level

For Review Only

(11, 12). As structural data is becoming more readily available, the wealth of information it provides should be incorporated into our understanding of the functional conservation of viral proteins throughout time. Indeed, recent studies have shown that structure-based phylogenetic analyses can outperform sequence-based phylogenetic analyses (9–17).

By converting the NSV NP sequence dataset into an AlphaFold2-derived structural dataset using flexible alignment, similar clustering patterns to the RdRp domain MSA emerged (**Figure 5B**). However, we did observe key differences. Of particular interest is the similarity between the NPs of the segmented *Serpentovirales* and the other segmented NSVs. While we did not estimate phylogenetic relationships using tree estimation algorithms, the above observation suggests that the current classification of the *Serpentovirales* may need to be critically examined. Electron microscopy analyses have shown that *serpentovirus* RNPs are flexible, having internally coiled loop-like structures, similar to the *bunyavirus* RNPs, and more linear collapsed duplex structures, similar to *articulavirus* RNPs (40). Morphologically, *serpentovirus* RNPs most closely resemble RNPs of viruses in the *tospoviridae* family (of the *bunyavirus* order), yet there is no evidence that serpentoviruses have enveloped virion particles and, as is the case for *tospoviruses*. Previous analysis of the overall architecture of NSV RNPs has shown a pattern in RNP flexibility that corresponds directly to genome organization (4). However, since there are only two micrographs available for *serpentovirales* RNPs, it is possible that this observation is an outlier rather than consistent across the seven species. Obtaining additional electron micrographs is needed to support a phylogenetic classification based on genome morphology and/or structure.

The above observations make it tempting to speculate about the evolutionary origin of NSV genome segmentation. The viral families that have been previously hypothesized to be associated with the evolution of NSV genome segmentation include *serpentoviruses* (within the *Ophioviridae*) along with viral species that belong to the *Chuviridae* (6). While all currently identified species of *Serpentovirales* have segmented genomes, *Chuviriales* with non-segmented and segmented genomes have been identified. Our current dataset includes 7 non-segmented *Chuvirales* that all belong to the *Jingchuvirales* order. These viruses firmly cluster with the nonsegmented NSVs in line with their genome organization. To obtain a better sense where the segmented *Chuvirales* would cluster, and potentially their position relative to the origin of NSV genome segmentation, our NP structural analysis can be extended in the future. Further analysis of the *Serpentovirales* may also provide greater insight into our understanding of the evolution of segmented NSV genomes.

It is possible that segmented NSV genomes evolved multiple times. This hypothesis could explain the large differences seen between articulaviruses and bunyaviruses in terms of the number of genome segments and the gene organization in the respective viral genomes. Moreover, several viral species that belong to the *monogenavirales* have segmented genomes, including orchid fleck dichorhavirus, which is classified within the *rhabdoviridae* and possesses two genome segments (6). Phylogenetic analysis suggests that this virus evolved from a non-segmented plant virus within the *rhabdoviridae* (39). Further analyses using larger datasets, multiple core genes, and tree reconstruction methods will likely be required to address these questions properly.

For Review Only

Lastly, we need to consider the likelihood that a multitude of viral species remains to be identified, and that the "origin species" of NSV genome segmentation has not been discovered yet. It is also possible that this species has been identified, but that it has so far been excluded from large scale evolutionary analyses because the RdRp sequences of this species were incomplete or missing. The findings presented here suggest a useful role for including structural information based on the sequences of non-RdRp proteins, such as NP, in phylogenomic analyses. We therefore hope that our findings will help inspire new research and ultimately the identification of a possible origin of NSV genome segmentation.

##### *Alignment analysis in R*

Original code to process and visually analyze the resulting alignment data was written in R. Heatmaps were created using R package gplots.

For Review Only

#### Data Availability and Requirements

Predicted AlphaFold structure files are available in pdb format in **Supplementary Files 1 and 2**. Dataset acquisition codes can be found in **Supplementary Table 1**.

#### Supplementary Information

Supplementary Table 1: Full NSV Dataset
Supplementary File 1: Initial Dataset NP PDBs (.pdb files)
Supplementary File 2: Full NSV Dataset NP AlphaFold PDBs (.pdb files)

#### Competing Interests

The authors declare no competing interests.

#### Acknowledgements

The authors would like to thank Dr. Ned Wingreen, Dr. Maria Rosenthal and Dr. Ingrida Olendraite for insightful discussions regarding the evolution of NSVs, as well as members of the Wingreen lab for discussion of the computational analyses. KRS was supported by NIH grants R01 GM140032 (to Ned Wingreen) and R01 AI170520 (to Adam Lauring, AJ te Velthuis and Colin Russell). ATV was supported by NIH grants DP2 AI175474 (to AJ te Velthuis) and R01 AI170520 (to Adam Lauring, AJ te Velthuis and Colin Russell), and Wellcome Trust and Royal Society grant 206579/17/Z (to AJ te Velthuis).

The authors are pleased to acknowledge that the work reported on in this paper was performed using the Princeton Research Computing resources at Princeton University which is a consortium of groups led by the Princeton Institute for Computational Science and Engineering (PICSciE) and Office of Information Technology's Research Computing.

ribonucleoprotein complex in negative-sense single-stranded RNA viruses. Trends Microbiol 21:475–484.

2. Luo M, Terrell JR, Mcmanus SA. 2020. Nucleocapsid Structure of Negative Strand RNA Virus. 8. Viruses 12:835.
3. Strauss JH, Strauss EG. 2008. CHAPTER 2 - The Structure of Viruses, p. 35–62. *In* Strauss, JH, Strauss, EG (eds.), Viruses and Human Disease (Second Edition). Academic Press, London.
4. Sabsay KR, Te Velthuis AJW. 2023. Negative and ambisense RNA virus ribonucleocapsids: more than protective armor. Microbiol Mol Biol Rev MMBR 87:e0008223.
5. Morin B, Kranzusch PJ, Rahmeh AA, Whelan SPJ. 2013. The polymerase of negative-stranded RNA viruses. Curr Opin Virol 3:103–110.
6. Li C-X, Shi M, Tian J-H, Lin X-D, Kang Y-J, Chen L-J, Qin X-C, Xu J, Holmes EC, Zhang Y-Z. 2015. Unprecedented genomic diversity of RNA viruses in arthropods reveals the ancestry of negative-sense RNA viruses. eLife 4:e05378.
7. STRAUSS JH, STRAUSS EG. 2008. Minus-Strand RNA Viruses. Viruses Hum Dis 137–191.
8. Wolf YI, Kazlauskas D, Iranzo J, Lucía-Sanz A, Kuhn JH, Krupovic M, Dolja VV, Koonin EV. 2018. Origins and Evolution of the Global RNA Virome. mBio 9:e02329-18.
9. Moi D, Bernard C, Steinegger M, Nevers Y, Langleib M, Dessimoz C. 2023. Structural phylogenetics unravels the evolutionary diversification of communication systems in gram-positive bacteria and their viruses. bioRxiv https://doi.org/10.1101/2023.09.19.558401.
10. Mifsud JCO, Lytras S, Oliver MR, Toon K, Costa VA, Holmes EC, Grove J. 2024. Mapping glycoprotein structure reveals defining events in the evolution of the Flaviviridae. bioRxiv https://doi.org/10.1101/2024.02.06.579159.
11. Krissinel E. 2007. On the relationship between sequence and structure similarities in

For Review Only

proteomics. Bioinformatics 23:717–723.

12. He Y, Maisuradze GG, Yin Y, Kachlishvili K, Rackovsky S, Scheraga HA. 2017. Sequence-, structure-, and dynamics-based comparisons of structurally homologous CheY-like proteins. Proc Natl Acad Sci 114:1578–1583.

13. Jumper J, Evans R, Pritzel A, Green T, Figurnov M, Ronneberger O, Tunyasuvunakool K, Bates R, Žídek A, Potapenko A, Bridgland A, Meyer C, Kohl SAA, Ballard AJ, Cowie A, Romera-Paredes B, Nikolov S, Jain R, Adler J, Back T, Petersen S, Reiman D, Clancy E, Zielinski M, Steinegger M, Pacholska M, Berghammer T, Bodenstein S, Silver D, Vinyals O, Senior AW, Kavukcuoglu K, Kohli P, Hassabis D. 2021. Highly accurate protein structure prediction with AlphaFold. 7873. Nature 596:583–589.

14. Mönttinen HAM, Ravantti JJ, Poranen MM. 2021. Structure Unveils Relationships between RNA Virus Polymerases. Viruses 13:313.

15. Malik AJ, Poole AM, Allison JR. 2020. Structural Phylogenetics with Confidence. Mol Biol Evol 37:2711–2726.

16. Sjölander K. 2010. Getting Started in Structural Phylogenomics. PLoS Comput Biol 6.

22. Turner PE. 2003. Searching for the advantages of virus sex. Orig Life Evol Biosphere J Int

For Review Only

Soc Study Orig Life 33:95–108.

28. Kuhn JH, Adkins S, Alkhovsky SV, Avšič-Županc T, Ayllón MA, Bahl J, Balkema-Buschmann A, Ballinger MJ, Bandte M, Beer M, Bejerman N, Bergeron É, Biedenkopf N, Bigarré L, Blair CD, Blasdell KR, Bradfute SB, Briese T, Brown PA, Bruggmann R, Buchholz UJ, Buchmeier MJ, Bukreyev A, Burt F, Büttner C, Calisher CH, Candresse T, Carson J, Casas I, Chandran K, Charrel RN, Chiaki Y, Crane A, Crane M, Dacheux L, Bó ED, de la Torre JC, de Lamballerie X, de Souza WM, de Swart RL, Dheilly NM, Di Paola N, Di Serio F, Dietzgen RG, Digiaro M, Drexler JF, Duprex WP, Dürrwald R, Easton AJ, Elbeaino T, Ergünay K, Feng G, Feuvrier C, Firth AE, Fooks AR, Formenty PBH, Freitas-Astúa J, Gago-Zachert S, García ML, García-Sastre A, Garrison AR, Godwin SE, Gonzalez J-PJ, de Bellocq JG, Griffiths A, Groschup MH, Günther S, Hammond J, Hepojoki J, Hierweger MM, Hongō S, Horie M, Horikawa H, Hughes HR, Hume AJ, Hyndman TH, Jiāng D, Jonson GB, Junglen S, Kadono F, Karlin DG, Klempa B,

For Review Only

Klingström J, Koch MC, Kondō H, Koonin EV, Krásová J, Krupovic M, Kubota K, Kuzmin IV, Laenen L, Lambert AJ, Lǐ J, Li J-M, Lieffrig F, Lukashevich IS, Luo D, Maes P, Marklewitz M, Marshall SH, Marzano S-YL, McCauley JW, Mirazimi A, Mohr PG, Moody NJG, Morita Y, Morrison RN, Mühlberger E, Naidu R, Natsuaki T, Navarro JA, Neriya Y, Netesov SV, Neumann G, Nowotny N, Ochoa-Corona FM, Palacios G, Pallandre L, Pallás V, Papa A, Paraskevopoulou S, Parrish CR, Pauvolid-Corrêa A, Pawęska JT, Pérez DR, Pfaff F, Plemper RK, Postler TS, Pozet F, Radoshitzky SR, Ramos-González PL, Rehanek M, Resende RO, Reyes CA, Romanowski V, Rubbenstroth D, Rubino L, Rumbou A, Runstadler JA, Rupp M, Sabanadzovic S, Sasaya T, Schmidt-Posthaus H, Schwemmle M, Seuberlich T, Sharpe SR, Shi M, Sironi M, Smither S, Song J-W, Spann KM, Spengler JR, Stenglein MD, Takada A, Tesh RB, Těšíková J, Thornburg NJ, Tischler ND, Tomitaka Y, Tomonaga K, Tordo N, Tsunekawa K, Turina M, Tzanetakis IE, Vaira AM, van den Hoogen B, Vanmechelen B, Vasilakis N, Verbeek M, von Bargen S, Wada J, Wahl V, Walker PJ, Whitfield AE, Williams JV, Wolf YI, Yamasaki J, Yanagisawa H, Ye G, Zhang Y-Z, Økland AL. 2022. 2022 taxonomic update of phylum Negarnaviricota (Riboviria: Orthornavirae), including the large orders Bunyavirales and Mononegavirales. Arch Virol 167:2857–2906.

29. Wang S, Ma J, Peng J, Xu J. 2013. Protein structure alignment beyond spatial proximity. 1. Sci Rep 3:1448.

30. Li SC. 2013. The difficulty of protein structure alignment under the RMSD. Algorithms Mol Biol AMB 8:1.

31. Li Z, Jaroszewski L, Iyer M, Sedova M, Godzik A. 2020. FATCAT 2.0: towards a better understanding of the structural diversity of proteins. Nucleic Acids Res 48:W60–W64.

32. Lennartz F, Hoenen T, Lehmann M, Groseth A, Garten W. 2013. The role of oligomerization for the biological functions of the arenavirus nucleoprotein. Arch Virol 158:1895–1905.

33. Turrell L, Hutchinson EC, Vreede FT, Fodor E. 2014. Regulation of Influenza A Virus

For Review Only

Nucleoprotein Oligomerization by Phosphorylation. J Virol 89:1452–1455.

34. Ariza A, Tanner SJ, Walter CT, Dent KC, Shepherd DA, Wu W, Matthews SV, Hiscox JA, Green TJ, Luo M, Elliott RM, Fooks AR, Ashcroft AE, Stonehouse NJ, Ranson NA, Barr JN, Edwards TA. 2013. Nucleocapsid protein structures from orthobunyaviruses reveal insight into ribonucleoprotein architecture and RNA polymerization. Nucleic Acids Res 41:5912–5926.
35. Zheng W, Tao YJ. 2013. Genome encapsidation by orthobunyavirus nucleoproteins. Proc Natl Acad Sci 110:8769–8770.
36. mattarnoldbio. 2023. AlphaPickle. Python.
37. García ML, Bó ED, da Graça JV, Gago-Zachert S, Hammond J, Moreno P, Natsuaki T, Pallás V, Navarro JA, Reyes CA, Luna GR, Sasaya T, Tzanetakis IE, Vaira AM, Verbeek M. 2017. ICTV Virus Taxonomy Profile: Ophioviridae. J Gen Virol 98:1161–1162.
38. Balique F, Lecoq H, Raoult D, Colson P. 2015. Can Plant Viruses Cross the Kingdom Border and Be Pathogenic to Humans? Viruses 7:2074–2098.
39. Jackson AO, Dietzgen RG, Goodin MM, Bragg JN, Deng M. 2005. Biology of Plant Rhabdoviruses. Annu Rev Phytopathol 43:623–660.
40. Ophioviridae | ICTV. https://ictv.global/report_9th/RNAneg/Ophioviridae. Retrieved 1 June 2023.
41. Madeira F, Pearce M, Tivey ARN, Basutkar P, Lee J, Edbali O, Madhusoodanan N, Kolesnikov A, Lopez R. 2022. Search and sequence analysis tools services from EMBL-EBI in 2022. Nucleic Acids Res 50:W276–W279.

For Review Only

| Number | Virus Abbreviation | Full Name | NCBI:txid |
|---|---|---|---|
| 1 | IAV | Influenza A Virus | 11320 |
| 2 | IBV | Influenza B Virus | 11520 |
| 3 | ICV | Influenza C Virus | 11552 |
| 4 | IDV | Influenza D Virus | 1511084 |
| 5 | ISAV | Infectious Salmon Anemia Virus, Salmon Isavirus | 55987 |
| 6 | JAQV | Johnston Atoll Quaranjavirus | 688437 |
| 7 | QQV | Quaranfil Quaranjavirus | 688436 |
| 8 | DhTV | Dhori Thogotovirus | 11318 |
| 9 | JoV | Jos Virus | 1027466 |
| 10 | BouV | Bourbon Virus | 1618189 |
| 11 | CCHRV | Crimean-Congo Hemorrhagic Fever Orthonairovirus | 1980519 |
| 12 | KUPEV | Kupe Virus | 498356 |
| 13 | HAZV | Hazara Virus | 11596 |
| 14 | ERVEV | Erve Virus | 248062 |
| 15 | JUNV | Junin Arenavirus, Argentinian mammarenavirus | 2169991 |
| 16 | TCRV | Tacaribe Virus, Tacaribe Mammarenavirus | 11631 |
| 17 | LFV | Lassa Fever Virus, Lassa Mammarenavirus | 11620 |
| 18 | BUNV | Bunyamwera Orthobunyavirus | 1933179 |
| 19 | LACV | La Crosse Virus | 11577 |
| 20 | LEAV | Leanyer Virus | 999729 |
| 21 | SBV | Schmallenberg Orthobunyavirus | 2560743 |
| 22 | TSWV | Tomato Spotted Wilt Virus, Tomato Spotted Wilt Orthotospovirus | 1933298 |
| 23 | RREV | Rose Rosette Emaravirus | 1980433 |
| 24 | ASAV | Ash Shoestring-Associated Emaravirus | 2854797 |
| 25 | PMEV | Perilla Mosaic Emaravirus | 2845802 |
| 26 | JSAV | Japanese Star Anise Ringspot-Associated Virus | 2798807 |
| 27 | ATOPV | Anopheles Triannulatus Orthophasmavirus | 2546222 |
| 28 | WMoV2 | Wuhan Mosquito Virus 2 | 1608127 |
| 29 | PGV2 | Pectinophora Gossypiella Virus 2 | 2856559 |
| 30 | SWSV2 | Sanxia Water Strider Virus 2 | 1608061 |

For Review Only

| | | | |
|---|---|---|---|
| 31 | SNV | Sin Nombre Orthohantavirus | 1980491 |
| 32 | ANDV | Andes Orthohantavirus | 1980456 |
| 33 | HTNV | Hantaan Orthohantavirus | 1980471 |
| 34 | RVF | Rift Valley Fever Virus | 11588 |
| 35 | TOSV | Toscana Virus | 11590 |
| 36 | GGV | Gouleako Goukovirus | 1980547 |
| 37 | LHQV1 | Linepithema Humile Qinvirus-like Virus 1 | 2259782 |
| 38 | FOAQV1 | Frankliniella Occidentalis Associated Qin-like Virus 1 | 2771469 |
| 39 | BlMaV | Blueberry Mosaic Associated Virus | 1520332 |
| 40 | CPsV | Citrus Psorosis Virus | 73561 |
| 41 | FSnV | Freesia Sneak Ophiovirus | 2170046 |
| 42 | LRNV | Lettuce Ring Necrosis Virus | 274495 |
| 43 | MLBvV | Mirafiore Lettuce Big-Vein Virus | 200255 |
| 44 | RWMV | Ranunculus White Mottle Virus | 152636 |
| 45 | TMmV | Tulip Mild Mottle Mosaic Virus | 219702 |
| 46 | WMV | Wuhan Mivirus | 2507319 |
| 47 | WTV2 | Wuhan Tick Virus 2 | 1608138 |
| 48 | ADMV | Amblyomma Dissimile Mivirus | 2875328 |
| 49 | BTV3 | Bole Tick Virus 3 | 1608042 |
| 50 | CTV2 | Changping Tick Virus 2 | 1608044 |
| 51 | STCV2 | Soybean Thrips Chu-like Virus 2 | 2800864 |
| 52 | NuoV | Nuomin Virus | 2916986 |
| 53 | VSV | Vesicular Stomatitis Virus | 11276 |
| 54 | GaV | Garba Virus | 864696 |
| 55 | OhV | Ohlsdorf Virus | 2040592 |
| 56 | BGV | Bahia Grande Virus | 932699 |
| 57 | MSV | Muir Springs Virus | 932700 |
| 58 | RABV | Rabies Lyssavirus | 11292 |
| 59 | DMSV | Drosophila Melanogaster Sigmavirus | 671497 |
| 60 | NDV | Newcastle Disease Virus, Avian Orthoavulavirus | 2560319 |
| 61 | HPIV5 | Human Parainfluenza Virus 5 | 2905673 |

For Review Only

| | | | |
|---|---|---|---|
| 62 | NiV | Nipah Virus | 121791 |
| 63 | CeMV | Cetacean Morbillivirus, Dolphin Morbillivirus | 36410 |
| 64 | MuV | Mumps Orthorubulavirus | 2560602 |
| 65 | MeV | Measles Morbillivirus | 11234 |
| 66 | SEN | Sendai Virus, Murine respirovirus | 11191 |
| 67 | RSV | Human Respiratory Syncytial Virus | 11250 |
| 68 | HyAV | Hymenopteran Arli-related Virus OKIAV100 | 2792564 |
| 69 | ABB1 | Aquatic Bird Bornavirus 1, Avian Bornavirus CG | 1715293 |
| 70 | BoDV | Borna Disease Virus | 12455 |
| 71 | NyNV | Nyamanini Nyavirus | 644610 |
| 72 | JPNV | Jeremy Point Nyavirus | 2652327 |
| 73 | FFV1 | Formica Fusca Virus 1 | 2018499 |
| 74 | WCV1 | Wenzhou Crab Virus 1 | 1608091 |
| 75 | OrV | Orinoco Virus | 1871345 |
| 76 | SJV | San Jacinto Virus | 2596788 |
| 77 | JCPV | Jungle Carpet Python Virus | 2016401 |
| 78 | SCPV | Southwest Carpet Python Virus, Southern Carpet Python Virus | 2016402 |
| 79 | SUDV | Sudan Ebolavirus | 186540 |
| 80 | EBOV | Ebola Virus, Zaire Ebola Virus | 128951 |

For Review Only

| Phylum | Genus | Family | Order |
|---|---|---|---|
| Polyploviricotina | Alpha influenza virus | Orthomyxoviridae | Articulavirales |
| Polyploviricotina | Beta influenza virus | Orthomyxoviridae | Articulavirales |
| Polyploviricotina | Gamma influenza virus | Orthomyxoviridae | Articulavirales |
| Polyploviricotina | Delta influenza virus | Orthomyxoviridae | Articulavirales |
| Polyploviricotina | Isavirus | Orthomyxoviridae | Articulavirales |
| Polyploviricotina | Quaranjavirus | Orthomyxoviridae | Articulavirales |
| Polyploviricotina | Quaranjavirus | Orthomyxoviridae | Articulavirales |
| Polyploviricotina | Thogotovirus | Orthomyxoviridae | Articulavirales |
| Polyploviricotina | Thogotovirus | Orthomyxoviridae | Articulavirales |
| Polyploviricotina | Thogotovirus | Orthomyxoviridae | Articulavirales |
| Polyploviricotina | Orthonairovirus | Nairoviridae | Bunyavirales |
| Polyploviricotina | Orthonairovirus | Nairoviridae | Bunyavirales |
| Polyploviricotina | Orthonairovirus | Nairoviridae | Bunyavirales |
| Polyploviricotina | Orthonairovirus | Nairoviridae | Bunyavirales |
| Polyploviricotina | Mammarenavirus | Arenaviridae | Bunyavirales |
| Polyploviricotina | Mammarenavirus | Arenaviridae | Bunyavirales |
| Polyploviricotina | Mammarenavirus | Arenaviridae | Bunyavirales |
| Polyploviricotina | Orthobunyavirus | Peribunyaviridae | Bunyavirales |
| Polyploviricotina | Orthobunyavirus | Peribunyaviridae | Bunyavirales |
| Polyploviricotina | Orthobunyavirus | Peribunyaviridae | Bunyavirales |
| Polyploviricotina | Orthobunyavirus | Peribunyaviridae | Bunyavirales |
| Polyploviricotina | Orthotospovirus | Tospoviridae | Bunyavirales |
| Polyploviricotina | Emaravirus | Fimoviridae | Bunyavirales |
| Polyploviricotina | Emaravirus | Fimoviridae | Bunyavirales |
| Polyploviricotina | Emaravirus | Fimoviridae | Bunyavirales |
| Polyploviricotina | Emaravirus | Fimoviridae | Bunyavirales |
| Polyploviricotina | Orthophasmavirus | Phasmaviridae | Bunyavirales |
| Polyploviricotina | Orthophasmavirus | Phasmaviridae | Bunyavirales |
| Polyploviricotina | Not classified | Phasmaviridae | Bunyavirales |
| Polyploviricotina | Sawastrivirus | Phasmaviridae | Bunyavirales |

For Review Only

| | | | |
|---|---|---|---|
| Polyploviricotina | Orthohantavirus | Hantaviridae | Bunyavirales |
| Polyploviricotina | Orthohantavirus | Hantaviridae | Bunyavirales |
| Polyploviricotina | Orthohantavirus | Hantaviridae | Bunyavirales |
| Polyploviricotina | Phlebovirus | Phenuiviridae | Bunyavirales |
| Polyploviricotina | Phlebovirus | Phenuiviridae | Bunyavirales |
| Polyploviricotina | Goukovirus | Phenuiviridae | Bunyavirales |
| Haploviricotina | Not classified | Qinviridae | Muvirales |
| Haploviricotina | Not classified | Qinviridae | Muvirales |
| Haploviricotina | Ophiovirus | Aspiviridae | Serpentovirales |
| Haploviricotina | Ophiovirus | Aspiviridae | Serpentovirales |
| Haploviricotina | Ophiovirus | Aspiviridae | Serpentovirales |
| Haploviricotina | Ophiovirus | Aspiviridae | Serpentovirales |
| Haploviricotina | Ophiovirus | Aspiviridae | Serpentovirales |
| Haploviricotina | Ophiovirus | Aspiviridae | Serpentovirales |
| Haploviricotina | Ophiovirus | Aspiviridae | Serpentovirales |
| Haploviricotina | Mivirus | Chuviridae | Jingchuvirales |
| Haploviricotina | Mivirus | Chuviridae | Jingchuvirales |
| Haploviricotina | Mivirus | Chuviridae | Jingchuvirales |
| Haploviricotina | Mivirus | Chuviridae | Jingchuvirales |
| Haploviricotina | Mivirus | Chuviridae | Jingchuvirales |
| Haploviricotina | Not classified | Chuviridae | Jingchuvirales |
| Haploviricotina | Not classified | Chuviridae | Jingchuvirales |
| Haploviricotina | Vesiculovirus | Rhabdoviridae | Mononegavirales |
| Haploviricotina | Sunrhavirus | Rhabdoviridae | Mononegavirales |
| Haploviricotina | Ohlsrhavirus | Rhabdoviridae | Mononegavirales |
| Haploviricotina | Barhavirus | Rhabdoviridae | Mononegavirales |
| Haploviricotina | Barhavirus | Rhabdoviridae | Mononegavirales |
| Haploviricotina | Lyssavirus | Rhabdoviridae | Mononegavirales |
| Haploviricotina | Sigmavirus | Rhabdoviridae | Mononegavirales |
| Haploviricotina | Orthoavulavirus | Paramyxoviridae | Mononegavirales |
| Haploviricotina | Orthorubulavirus | Paramyxoviridae | Mononegavirales |

| | | | |
|---|---|---|---|
| Haploviricotina | Henipavirus | Paramyxoviridae | Mononegavirales |
| Haploviricotina | Morbillivirus | Paramyxoviridae | Mononegavirales |
| Haploviricotina | Orthorubulavirus | Paramyxoviridae | Mononegavirales |
| Haploviricotina | Morbillivirus | Paramyxoviridae | Mononegavirales |
| Haploviricotina | Respirovirus | Paramyxoviridae | Mononegavirales |
| Haploviricotina | Orthopneumovirus | Pneumoviridae | Mononegavirales |
| Haploviricotina | Arlivirus | Lispiviridae | Mononegavirales |
| Haploviricotina | Orthobornavirus | Bornaviridae | Mononegavirales |
| Haploviricotina | Orthobornavirus | Bornaviridae | Mononegavirales |
| Haploviricotina | Nyavirus | Nyamiviridae | Mononegavirales |
| Haploviricotina | Nyavirus | Nyamiviridae | Mononegavirales |
| Haploviricotina | Formivirus | Nyamiviridae | Mononegavirales |
| Haploviricotina | Crustavirus | Nyamiviridae | Mononegavirales |
| Haploviricotina | Orinovirus | Nyamiviridae | Mononegavirales |
| Haploviricotina | Nyavirus | Nyamiviridae | Mononegavirales |
| Haploviricotina | Carbovirus | Bornaviridae | Mononegavirales |
| Haploviricotina | Carbovirus | Bornaviridae | Mononegavirales |
| Haploviricotina | Ebolavirus | Filoviridae | Mononegavirales |
| Haploviricotina | Ebolavirus | Filoviridae | Mononegavirales |

For Review Only

| RdRp Sequence Accession | NP Sequence Accession |
|---|---|
| PB1: NC_007358.1:25-2298, PB2: NC_002023.1:28-2307, PA: NC_002022.1:25-21 | BBB04703.1 |
| PB1: NC_002204.1:21-2279, PB2: NC_002205.1, PA: NC_002206.1:1-2181 | ABN50464.1 |
| PB1: NC_006308.2:18-2282, PB2: NC_006307.2:22-2346, PA: NC_006309.2:22-21 | BAV18699.1 |
| PB1: NC_036615.1:26-2287, PB2: NC_036616.1:15-2333, PA: NC_036619.1:23-21 | YP_009449558.1 |
| PB1: NC_006503.1:35-2161, PB2: NC_006505.1, PA: NC_006501.1:5-1741 | AAM12950.1 |
| PB1: NC_052931.1:34-2367, PB2: NC_052925.1:28-2379, PA: NC_052926.1:26-23 | YP_009996581.1 |
| PB1: NC_038821.1:33-2366, PB2: NC_038817.1:c2391-190, PA: NC_038818.1:29- | AXL67890.1 |
| PB1: NC_034263.1:25-2175, PB2: NC_034261.1:52-2334, PA: NC_034254.1:29-19 | YP_009352881.1 |
| PB1: HM627170.1, PB2: HM627174.1, PA: HM627175.1 | AED98374.1 |
| PB1: KU708254.2, PB2: KU708253.2, PA: MK453527.1 | QCO69323.1 |
| NC_005301.3:77-11914 | AZS18999.1 |
| EU257628.1 | pdb\|4XZE\|A Chain A |
| NC_038709.1:38-11809 | sp\|P27318.1\|NCAP_HAZVJ |
| JF911697.1 | AFH89034.1 |
| NC_005080.1:c7084-452 | AVD30030.1 |
| NC_004292.1:c7072-440 | AGC92408.1 |
| NC_004297.1:c7122-466 | AAG41803.1 |
| NC_001925.1:51-6767 | AAL37356.1 |
| NC_004108.1:62-6853 | NP_671970.1 |
| NC_043563.1:39-6821 | pdb\|4J1J\|C Chain C |
| NC_043583.1:28-6792 | BCH47625.1 |
| NC_002052.1:34-8661 | BAB18309.1 |
| NC_015298.1:c6919-89 | UME38974.1 |
| OU466880.1 | CAG9003605.1 |
| LC721296.1 | BDH47810.1 |
| LC597437.1 | BCO17110.1 |
| NC_055394.1:91-6429 | YP_010086187.1 |
| NC_031312.1:151-6486 | YP_009305134.1 |
| QID77675.1 | QXL90808.1 |
| NC_055192.1:55-7089 | YP_010085073.1 |

For Review Only

| | |
|---|---|
| NC_005217.1:36-6497 | pdb\|5E06\|A Chain A |
| NC_003468.2:36-6497 | QRY27104.1 |
| NC_005222.1:38-6493 | ALI59823.1 |
| NC_014397.1:19-6297 | ABP88854.1 |
| MT032308.1 | AIQ84589.1 |
| NC_043051.1:19-6234 | YP_009664620.1 |
| MH213241.1 | AXA52553.1 |
| NS31049.1 | QNS31050.1 |
| NC_036635.1:c7252-239 | YP_009449564.1 |
| NC_006314.1:c7420-170 | AAC41022.1 |
| *Not available* | AWJ64344.1 |
| NC_006051.1:c6958-125 | QTP72399.1 |
| NC_011558.1:339-6461 | AAU12876.1 |
| NC_043389.1:c4018-1 | AAT08132.1 |
| *Not available* | AWJ64310.1 |
| MZ244267.1:67-6636 | QYW06803.1 |
| NC_028266.1:72-6641 | UGM46212.1 |
| MZ502309.1:98-6655 | UAJ23572.1 |
| NC_028259.1:77-6544 | QYW06797.1 |
| NC_028260.1:67-6537 | AJG39046.1 |
| QQP18758.1 | QQP18725.1 |
| MW029967.1:86-6601 | UKS70442.1 |
| X00939.1 | AAA48470.1 |
| NC_055530.1:4599-10817 | YP_010087296.1 |
| NC_055477.1:4977-11459 | YP_010086782.1 |
| NC_055531.1:5868-12395 | YP_010087303.1 |
| NC_055532.1:5859-12386 | YP_010087309.1 |
| NC_001542.1:5388-11863 | ARM19989.1 |
| NC_013135.1:6000-12389 | ACV67016.1 |
| NC_039223.1:8387-15001 | BAO02663.1 |
| KY114804.1:8414-15181 | AAA47880.1 |

For Review Only

| | |
|---|---|
| NC_002728.1:11259-18213 | AAX51852.1 |
| NC_005283.1:9042-15593 | AYR16894.1 |
| NC_002200.1:8438-15223 | QLH93297.1 |
| NC_001498.1:9212-15854 | CAC34604.1 |
| NC_001552.1:8556-15242 | AAB06278.1 |
| NC_001803.1:8460-15037 | AIY60635.1 |
| MW288216.1:5122-11216 | QPL15361.1 |
| NC_029642.1:2398-8827 | UGC11957.1 |
| NC_001607.1:3687-8822 | QJI08281.1 |
| NC_012703.1:c5877-67 | YP_002905342.1 |
| MN045233.1:6858-12659 | QFG01726.1 |
| MH477287.1:4098-9731 | AYW51534.1 |
| NC_031275.1:4119-9542 | YP_009304556.1 |
| NC_043485.1:c5293-1 | YP_009666287.1 |
| MK971153.1:7383-13214 | QFQ60714.1 |
| NC_039013.1:3571-8754 | YP_009508479.1 |
| NC_039014.1:3556-8733 | YP_009508485.1 |
| NC_006432.1:11457-18494 | AWK96640.1 |
| NC_002549.1:11501-18282 | CAA70541.1 |

For Review Only
