## Supplemental Information for "Using structure prediction of negative sense RNA virus nucleoproteins to assess evolutionary relationships"

**Data exclusion**
Overview of NPs that were folded with AlphaFold2, but excluded from the dataset due to low confidence scores (average PAE scores <15 and average pLDDT scores >80).

1. Genoa virus
2. Sichuan mosquito mivirus
3. Culex mosquito virus 4
4. Hardyhead chuvirus
5. Soybean cyst nematode socyvirus
6. Bremia lactucae associated yuevirus-like Virus 1
7. Wuham millipede virus 2
8. Fushun phasmavirus 1
9. Sanya sesamia inferens phasmavirus 1
10. Wuchange cockroach virus 1
11. Wuhan insect virus 2
12. Pear chlorotic leaf spot-associated virus
13. Tulasnella bunyavirales-like virus 1
14. Wenzhou crab virus 2
15. Wuhan louse fly virus 7
16. Turtle fraservirus 1
17. Chatham eel tosovirus

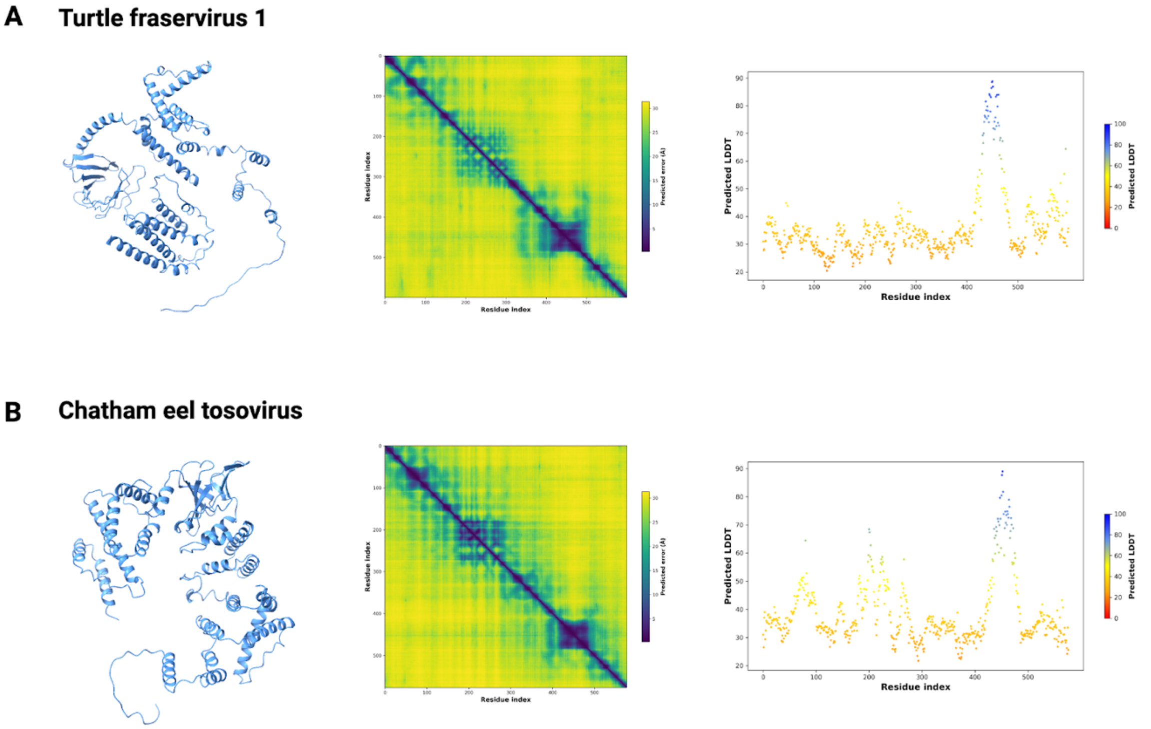


**Supplemental Figure 1: AlphaFold2 Predictions of NPs encoded by *Tosoviridae* that we excluded from the clustering analysis*.*** Structure predictions of (**A**) urtle fraservirus 1 (TFV1) and (**B**) Chatham eel tosovirus (CETV) based on available amino acid sequences UJT32109.1 and WLJ60756.1, respectively. The highest quality model prediction is shown on the left in blue ribbon format. The predicted alignment error (PAE) is shown as a heatmap in the middle. High confidence predictions are those with an average PAE of less than 15 angstroms. The calculated average PAE scores are 26.67 and 25.83 angstroms for TFV1 and CETV, respectively. The predicted per-residue model confidence score (pLDDT score; range of 1-100) is plotted over all residues for each model on the right. A high confidence prediction has an average pLDDT score of greater than 80. The calculated average pLDDT scores are 36.15 and 38.77 for TFV1 and CETV, respectively. The region of highest confidence in the turtle fraservirus 1 nucleoprotein corresponds to the peak seen in the pLDDT plot from residues 431 to 463 which has an average pLDDT of 78 and an average PAE of 2.5 angstroms. This corresponds to a region whose sequence aligns only to itself (Blastp) and is structurally composed of an alpha helix and tail, which is too small/vague of a structure to attempt to structurally align to any other nucleoprotein and expect significant results.

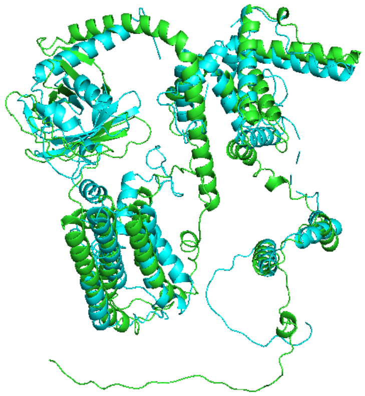

**Supplemental Figure 2: FATCAT structure alignment of AlphaFold2 predictions of TFV1 and CETV nucleoproteins.** Alignment of the two predicted structure models of TFV1 (green) and CETV (blue) NPs shows significant alignment with the allowance of 5 twists and 6 blocks. Structurally aligning the entire predicted structures of TFV1 and CETV using the FATCAT algorithm yields a significant p-value of 7.7E-04, suggesting that these (low-confidence predictions) are highly similar structures, as expected given that they are both from Tosoviridae (FIGURE). By sequence, the TFV1 and CETV nucleoproteins are 40.4% identical (blastp). Unlike the AlphaFold predictions for EBOV and SUDV, the low confidence for TFV1 and CETV cannot be explained by a high proportion of intrinsic disorder (calculated by IUPred2) with less than 20% of the structure predicted to be intrinsically disordered (10.2% for TFV1 and 3.0% for CETV). The low confidence is likely a result of the lack of data available for this viral family and the dissimilarity to other known viral nucleoproteins, which has consequently prevented Tosoviridae from being classified into an order, class, or subphylum by traditional phylogenomic methods. Visually assessing the NP models for TFV1 and CETV show that the regions of higher structure, corresponding to the darker blue clusters on the PAE heatmaps are very loosely connected. Both of these structures appear to be loose with high predicted aligned error between the individual regions of secondary structure. Structural alignment to several NSV NPs (IAV, BUNV, MLBvV, EBOV) produces insignificant results with p-values of 0.148, 0.397, 0.418, 0.256, respectively.
